# Diving into bacterial dormancy: emergence of osmotically stable wall-less forms in an aquatic environment

**DOI:** 10.1101/2023.11.16.566987

**Authors:** Filipe Carvalho, Alexis Carreaux, Anna Sartori-Rupp, Stéphane Tachon, Anastasia D. Gazi, Pascal Courtin, Pierre Nicolas, Florence Dubois-Brissonnet, Aurélien Barbotin, Emma Desgranges, Karine Gloux, Catherine Schouler, Rut Carballido-López, Marie-Pierre Chapot-Chartier, Eliane Milohanic, Hélène Bierne, Alessandro Pagliuso

## Abstract

Bacteria can respond to environmental stresses by entering a dormant state, called viable but non-culturable (VBNC) state, in which they no longer grow in routine culture media. VBNC pathogens pose thus a significant risk for human and animal health as they are not detected by standard growth-based techniques and can “wake up” back into a vegetative and virulent state. Although hundreds of species were reported to become VBNC in response to different stresses, the molecular mechanisms governing this phenotypic switch remain largely elusive.

Here, we characterized the VBNC state transition process in the Gram-positive pathogen *Listeria monocytogenes* in response to nutritional deprivation. By combining fluorescence microscopy, cryo-electron tomography and analytical biochemistry, we found that starvation in mineral water drives *L. monocytogenes* into a VBNC state via a mechanism of cell wall (CW) shedding that generates osmotically stable CW-deficient (CWD) coccoid forms. This phenomenon occurs in multiple *L. monocytogenes* strains and in other *Listeria* species, suggesting it may be a stress-adapting process transversal to the *Listeria* genus. Transcriptomic and gene-targeted approaches revealed the stress response regulator SigB and the autolysin NamA as major moderators of CW loss and VBNC state transition. Finally, we show that this CWD dormant state is transient as VBNC *Listeria* revert back to a walled, vegetative and virulent state after passage in embryonated eggs.

Our findings provide unprecedented detail on the mechanisms governing the transition to a VBNC state, and reveal that dormant CWD bacterial forms can naturally arise in aquatic environments without osmotic stabilization. This may represent an alternative strategy for bacterial survival in oligotrophic conditions, which can potentially generate public health-threatening reservoirs of undetectable pathogens.

## Introduction

Bacteria often face less than optimal growth conditions and a variety of abiotic stresses in their environment. Some species are able to produce highly resistant cellular structures, called endospores, to enter a metabolically inactive state until environmental conditions are adequate for resuming vegetative growth (Beskrovnaya et al., 2021). Alternatively, bacteria may enter a dormant state known as the viable but non-culturable (VBNC) state, in which they preserve some metabolic activity at the expense of losing the ability to grow on regular culture media (Dong et al., 2020). The VBNC state is documented in over a hundred species (Ayrapetyan et al., 2018; Dong et al., 2020), but our knowledge concerning the molecular processes driving the transition from a vegetative lifestyle to the VBNC state, particularly in Gram-positive bacteria, are still fragmentary.

The transition to a VBNC state is frequently accompanied by a morphological change, often cell dwarfing and/or rounding (Li et al., 2014). The underlying reasons are not entirely understood but an hypothesis is that a spherical shape, with a smaller surface area/volume ratio, might help VBNC bacteria reduce their energy demands and optimize nutrient uptake (Baker et al., 1983). Studies mostly performed in Gram-negative bacteria reported structural modifications of the cell wall (CW) peptidoglycan recovered from VBNC cells that might contribute to cell rounding (Costa et al., 1999; Signoretto et al., 2000, 2002). However, a direct link between CW modifications, cell morphology and dormancy has not yet been clearly established.

Cell rounding has also been observed when some bacteria switch into a CW-deficient (CWD) state. This is the case of L-forms, CWD variants generated by exposure to CW-targeting agents, such as wall-active antibiotics, lytic enzymes or phages (Errington et al., 2016; Kawai et al., 2018; Wohlfarth et al., 2023). L-form cells remain viable and able to replicate, but the absence of CW makes them sensitive to osmotic lysis, and therefore they need to be cultivated in osmoprotective conditions (Errington et al., 2016). For this reason, the physiological relevance of L-forms, and CWD bacteria in general, is a matter of debate (Errington et al., 2016).

Here, we report that the Gram-positive bacterium *Listeria monocytogenes* (*Lm*) undergoes a rod-to-coccus differentiation in transition to a VBNC state in a nutrient-deprived natural water setting. We reveal that this cell rounding results from loss of the CW via a molting-like shedding process. Remarkably, these CWD VBNC *Lm* forms are resistant to osmotic lysis, likely as a result of adaptive changes in the physicochemical properties of their plasma membrane. We further show that this CWD VBNC state is extensive to other *Listeria sensu stricto* species. To our knowledge, this is the first report of CWD VBNC bacteria naturally arising in a non-osmotically stabilized environment. We further identified the stress-responsive transcription factor SigB and the autolysin NamA as major molecular players in the formation of CWD VBNC *Lm*. Finally, we show that dormant wall-less *Lm* can revert back to a walled, vegetative and fully virulent state after passage in embryonated chicken eggs. Our results suggest that CW shedding is an adaptive process employed by *Listeria* to survive under prolonged nutritional limitation.

## Results

### Dynamics of VBNC *Lm* formation in mineral water

To induce a VBNC state in *Lm*, we followed a starvation-based approach by incubating *Lm* in water (Besnard et al., 2000b, 2002). We used a commercial mineral water due to its natural spring origin, low mineral content and quality-controlled composition (**Supplementary Table 1**). As the starting number of bacteria affects the dynamics of culturability loss (Besnard et al., 2002), we tested initial *Lm* concentrations ranging from 10^9^ to 10^6^ bacteria/mL. We observed that the rate and magnitude of culturability loss increased when the starting bacterial concentration was reduced (**Fig. 1A**). Notably, a concentration of 10^6^ *Lm*/mL resulted in culturability levels of <1 colony-forming units (CFU)/mL after 28 days (**Fig. 1A**). We chose a starting concentration of 10^8^ *Lm*/mL as the standard condition to induce the VBNC state in mineral water throughout this work, since it produces a 2-log drop in culturable *Lm* after 28 days while still providing sufficient material for downstream analyses.

**Fig. 1.**
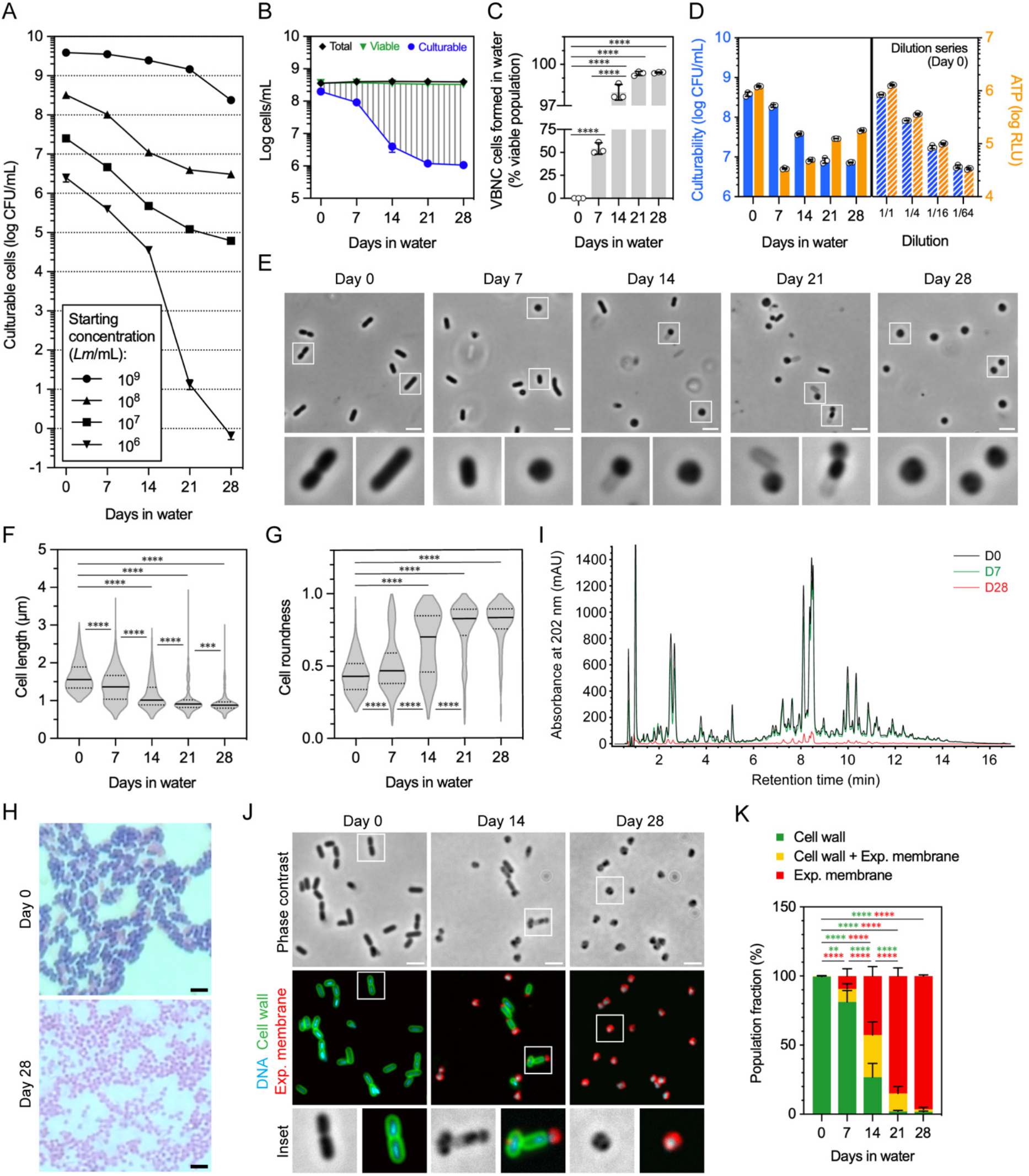
*Lm* transitions into a CWD VBNC state after prolonged incubation in mineral water. **(A)** Culturability profiles of *Lm* EGDe in mineral water suspensions with different starting bacterial concentrations. The number of culturable bacteria was determined at each timepoint by enumeration of colony-forming units (CFUs). Data are represented as the mean ± standard deviation (SD) from three independent suspensions. **(B)** Total, viable and culturable cell profiles of *Lm* EGDe in mineral water suspensions (10^8^ cells/mL). Culturable bacteria were determined by enumeration of CFUs, while total and viable bacteria were quantified by flow cytometry using CFDA. Data are represented as the mean ± SD from three independent suspensions. The area covered with vertical grey dashes indicates the VBNC cell population. **(C)** Fraction of the bacterial population consisting of VBNC cells formed in mineral water. Data are represented as the mean ± SD from three independent suspensions, each indicated by a dot. Statistical analysis was performed using one-way ANOVA with Tukey’s post hoc multiple comparison correction. ****, p≤0.0001. **(D)** Comparison of the culturability and ATP content of *Lm* EGDe sampled from mineral water suspensions at the indicated timepoints. Culturable bacteria were determined by enumeration of CFUs, while ATP levels were quantified with a luciferase-based assay. The graph on the right shows the culturability and ATP levels assessed from serial dilutions of *Lm* EGDe suspensions, immediately after preparation (day 0), and serves as a reference for the ATP content expected from a defined number of culturable cells. Data are represented as the mean ± SD from three independent suspensions, each indicated by a dot. **(E)** Phase-contrast micrographs of *Lm* EGDe sampled from mineral water suspensions at the indicated timepoints. Bacteria highlighted by white squares are shown enlarged in bottom panels. Scale bar: 2 µm. **(F, G)** Quantification of the cell length (F) and roundness (G) of *Lm* EGDe sampled from mineral water suspensions at the indicated timepoints. Data are represented as violin plots with the median (solid line) + interquartile range (dashed lines) from three independent suspensions (n=800–1600 cells per timepoint). Statistical analysis was performed using a Kruskal-Wallis test with Dunn’s post hoc multiple comparison correction. ****, p≤0.0001. **(H)** Micrographs of Gram-stained *Lm* EGDe sampled from mineral water suspensions at the beginning (day 0) and end (day 28) of the incubation period. Scale bar: 2 µm. **(I)** Reverse-phase UHPLC muropeptide profile obtained from peptidoglycan extracted from *Lm* EGDe suspensions in mineral water on the first day (D0) and after 7 (D7) and 28 (D28) days. The amount of material analyzed was normalized to the same bacterial cell number (1.5×10^9^). Profiles are representative of two independent suspensions. **(J)** Phase-contrast and fluorescence micrographs of *Lm* EGDe sampled from mineral water suspensions (10^8^ cells/mL) at the indicated timepoints. Bacteria were fixed and fluorescently labelled for visualization of DNA (Hoechst 33342), CW (WGA) and exposed plasma membrane (anti-CWD *Lm*). Bacteria highlighted by white squares are shown enlarged in the bottom panels. Scale bar: 2 µm. **(K)** Fraction of *Lm* EGDe in mineral water suspensions showing single- or double-labeling of CW (WGA) and exposed plasma membrane (anti-CWD *Lm*) by fluorescence microscopy at the indicated timepoints. Data are represented as stacked bars (one for each labelling group) with mean ± SD from three independent suspensions. Statistical analysis was performed using a two-way ANOVA test with Tukey’s post hoc multiple comparison correction, and only indicated for the single-labelled groups. **, p≤0.01; ****, p≤0.0001.

To confirm the formation of VBNC *Lm*, we monitored the total number of bacteria and the fraction of viable bacteria by flow cytometry. The viable population was determined using carboxyfluorescein diacetate (CFDA, **Extended Data Fig. 1A**), a fluorogenic dye that is enzymatically activated and retained in the cytoplasm of metabolically active bacteria with an integral plasma membrane (Wideman et al., 2021). While CFU counts dropped progressively to 10^6^ *Lm*/mL after 28 days, the total and viable population numbers remained nearly unchanged (**Fig. 1B**). The increasing difference between the viable and culturable populations with time demonstrates the gradual and almost complete transition to a VBNC state (**Fig. 1C**). This was also observed when using the double-dye Live/Dead assay to assess viability (**Extended Data Fig. 1B**). In this case, viable population numbers slightly dropped with time, which might be related with the different mode of action of these dyes (Stiefel et al., 2015).

ATP is only produced by live cells and quickly depleted upon cell death, constituting thus a marker of cellular viability. We measured the intracellular ATP levels in mineral water-incubated *Lm* cells over time, in parallel to their culturability. Variations in the ATP content did not follow the changes in culturability, unlike what we observed from a dilution series of freshly prepared suspensions, used to report the ATP content expected from a given number of culturable cells (**Fig. 1D**). Indeed, whereas culturable *Lm* numbers declined steadily, ATP levels dropped drastically after 7 days, recovering partially afterwards. Importantly, from day 21, the measured ATP levels were higher than those expected from a similar number of culturable cells (**Fig. 1D**), suggesting that this ATP surplus comes from the larger, non-culturable *Lm* subpopulation.

Together, these results confirm the efficient transition of *Lm* to a VBNC state in mineral water.

### VBNC *Lm* assume a coccoid morphology in mineral water

Changes in the bacterial cell size and shape are frequently associated with the VBNC state (Li et al., 2014). We thus acquired phase-contrast images of *Lm* suspensions in mineral water to track the occurrence of morphological changes during VBNC cell formation. The initial *Lm* population (day 0) consisted of typical rod-shaped cells, isolated or in tethered pairs (**Fig. 1E**). From 7 days of incubation, coccoid forms were also observed and became more abundant over time, at the expense of the rod-shaped subpopulation (**Fig. 1E**). Quantitative analysis of the populational morphology confirmed this progressive rod-to-coccus shape transformation (**Fig. 1F, G**). A suspension of GFP-expressing *Lm* showed GFP-positive coccoid cells appearing as of day 7 and increasing in number by day 28 (**Extended Data Fig. 1**), further confirming that the spherical forms derive from the initial rod-shaped *Lm*. Noteworthy, coccoid cells were sometimes found next to phase-light rod-shaped structures resembling empty cell wall sacculi; in some cases, even appearing to extrude from the latter (**Fig. 1E**, insets day 14 and 21).

These results show that incubation in mineral water triggers a rod-to-coccus transition in *Lm* cells. Interestingly, the dynamics of this morphological transition closely overlap with the dynamics of culturability decline and VBNC cell formation (**Fig. 1B, C**), suggesting a link between the shape change and the transition to a VBNC state.

### Rod-to-coccus transition in *Lm* is caused by CW loss via a molting-like shedding process

The switch from rod to coccoid shape has been observed in bacteria converting to L-forms, with the loss of the CW as the main driving force of this morphological change (Dell’Era et al., 2009; Domínguez-Cuevas et al., 2012; Errington et al., 2016). Having observed coccoid *Lm* associated with “ghost” structures resembling empty cell wall sacculi (**Fig. 1E**), we investigated whether coccoid *Lm* cells were CWD forms. We first performed a Gram staining of the *Lm* population at day 0 and day 28. Remarkably, the usual crystal violet staining displayed by rod-shaped *Lm* cells at day 0 was no longer present when the population consisted of coccoid cells after 28 days in mineral water (**Fig. 1H**), indicating the absence of a typical Gram-positive CW in coccoid VBNC *Lm* cells. We then compared the peptidoglycan content purified from equivalent *Lm* cell numbers at different timepoints of incubation in water, by performing UHPLC analysis of muropeptides. The muropeptide elution profiles showed that, while their composition did not visibly change, the amount of the different muropeptide species decreased with time until virtually no peptidoglycan was detected by day 28 (**Fig. 1I**). Together, these results confirm the progressive depletion of the *Lm* CW during transition to a VBNC state.

Next, we monitored the dynamics of *Lm* CW loss by fluorescence microscopy. The walled *Lm* population was fluorescently stained with wheat germ agglutinin (WGA), a lectin that binds to free *N*-acetylglucosamine (GlcNAc) residues present in wall teichoic acids (WTAs) of serogroup 1/2 strains (Fiedler, 1988). To monitor in parallel the appearance of CWD *Lm* cells, we generated anti-CWD *Lm* antibodies after immunization of rabbits with CWD *Lm* recovered from mineral water suspensions after 28 days. The anti-CWD *Lm* antibodies labeled the coccoid (i.e. CWD) but not the rod-shaped (i.e. walled) *Lm* cells, confirming their specificity (**Extended Data Fig. 1**). On the first day, *Lm* cells were only stained by WGA, indicating intact CW and a plasma membrane externally inaccessible to labeling by the anti-CWD *Lm* (**Fig. 1J, K**). Two additional subpopulations emerged with time: double-labelled cells, corresponding to *Lm* with a more permeable/damaged CW, and cells labeled only by the anti-CWD *Lm*, representing CWD *Lm* (**Fig. 1J, K**). The fraction of CWD *Lm* progressively increased with time to represent >95% of bacteria by day 28 (**Fig. 1K**).

Lastly, we investigated the CW loss phenomenon at the ultrastructural level in near-native conditions by cryogenic electron tomography (cryo-ET) of whole *Lm* cells. Micrographs and 3D rendering of segmented tomograms allowed us to reconstruct the different stages of the morphological transition of *Lm* in mineral water. Starting as a rod-shaped bacterium with a CW wrapped tightly around the plasma membrane (**Fig. 2, stage 0; Supplementary Movie 1**), *Lm* cells start showing a substantial detachment between these two layers (**Fig. 2, stage 1**), followed by the weakening and appearance of variably sized gaps in the CW mesh (**Fig. 2, stage 2; Supplementary Movie 2**). These gaps allow the enclosed protoplast to gradually egress the CW sacculus (**Fig. 2, stages 3 and 4; Supplementary Movies 3 and 4**) and escape into the extracellular medium as a spherical cell (**Fig. 2, stage 5; Supplementary Movie 5**).

**Fig. 2.**
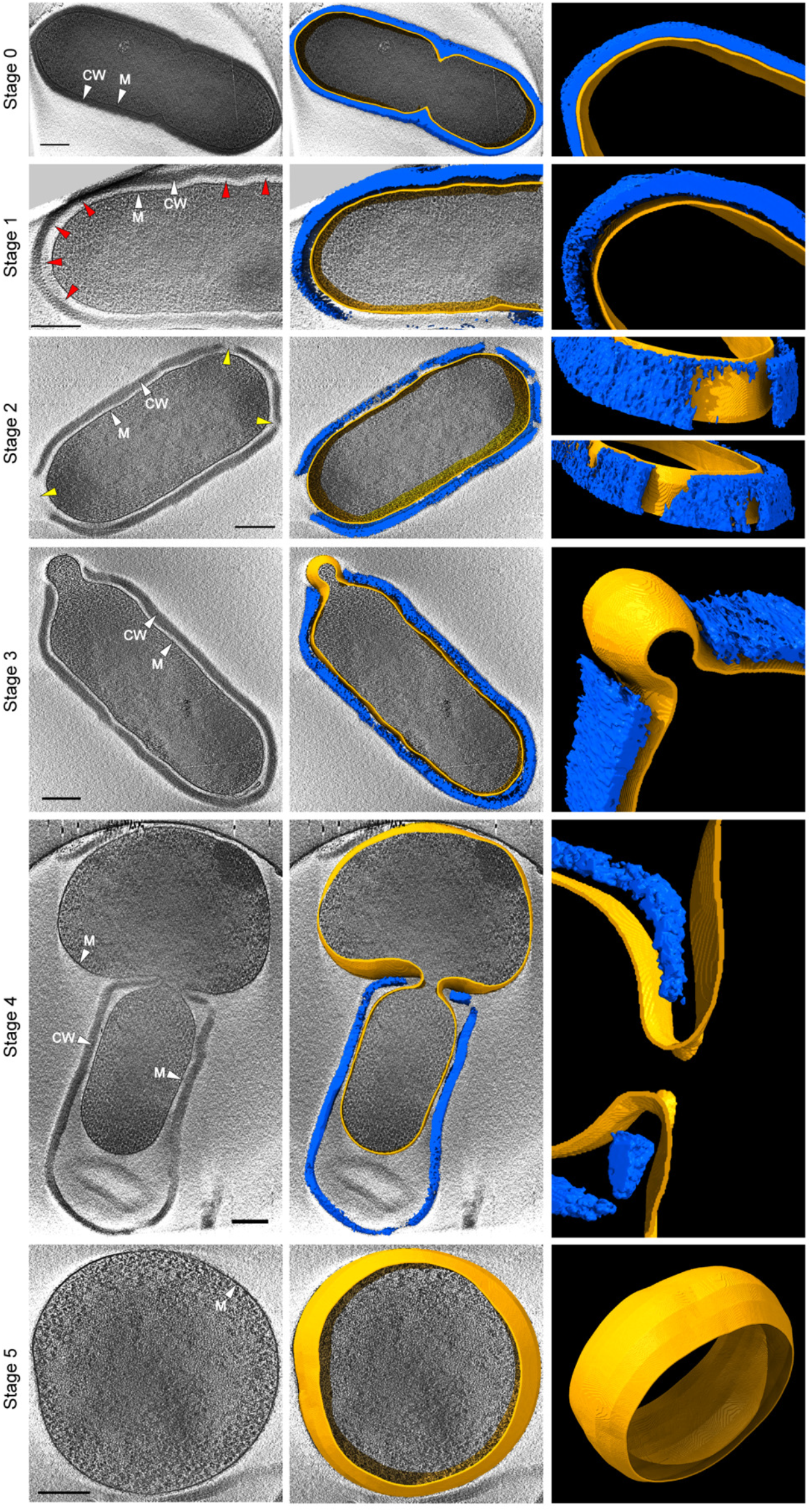
Cryo-electron tomography of the *Lm* CW shedding stages during transition to the VBNC state. Ultrastructure of the different *Lm* CW shedding stages obtained by cryo-electron tomography of *Lm* EGDe sampled from mineral water suspensions (10^8^ *Lm*/mL) between day 0 and day 14. **Stage 0**: rod-shaped bacterium with tightly connected plasma membrane and CW. In this example, a septum is forming at the midcell. **Stage 1**: the detachment of the plasma membrane and CW layers results in the formation of a periplasm-like space. **Stage 2**: breaches of variable size are formed in the CW matrix, exposing the enclosed protoplast to the outside environment. **Stage 3**: the bacterial protoplast begins bulging through a CW breach at one of the poles. **Stage 4**: the protoplast continues squeezing out, leaving an empty cell wall sacculus behind. **Stage 5**: the bacterium has fully molted from its rod-shaping cell wall, assuming a coccoid morphology as a CWD form. Left panels display a tomogram slice, in which CW and membrane (MB) are indicated by white arrowheads, a perisplam-like space is indicated by red arrowheads, and CW breaches are indicated by yellow arrowheads. Right panels display a 3D rendering of the CW (blue) and membrane (orange), obtained by manual segmentation of tomogram slices, and middle panels display a superposition of the tomogram slice and the 3D models. Scale bar: 200 nm. Movies showing all the tomogram slices used for 3D reconstruction of the bacterial CW and membrane for stages 0 and 2–5 are available as supplementary Movies S1 to S4.

Altogether, these results reveal that *Lm* cells transitioning to a VBNC state lose their CW through a molting-like shedding process that generates wall-less coccoid cell forms.

### A CWD VBNC state is widespread in *Listeria* species

We next wondered if the CWD VBNC state induced in mineral water occurred in other *Lm* strains, besides our reference strain EGDe. We monitored VBNC cell formation in the *Lm* strain 10403S, another well-studied reference laboratory strain (Bishop & Hinrichs, 1987), and in two clinical *Lm* strains isolated from human and bovine listeriosis cases: CLIP 63713 and JF5203, respectively (Aguilar-Bultet et al., 2018; Jonquières et al., 1998). Similar to EGDe, these three strains showed declining culturability over time (**Extended Data Fig. 1A–D**) and formation of a VBNC subpopulation (**Extended Data Fig. 1E–H**) in mineral water. To monitor the presence of CW by fluorescence microscopy, we used a commercial antibody raised against *Lm* CW-specific antigens (anti-*Lm*), since WGA does not label the CW of *Lm* serogroup 4 strains (CLIP 63713 and JF5203) due to the lack of free GlcNAc residues in their WTAs (Rismondo et al., 2020; Shen et al., 2017). The three strains displayed gradual loss of the CW (**Extended Data Fig. 1I–L, Q**) and exposure of the plasma membrane (**Extended Data Fig. 1M–P, Q**), with dynamics comparable to those of VBNC cell formation. These results demonstrate that the formation of CWD VBNC forms is a strain-independent property of *Lm*.

We then investigated other *Listeria* species, namely from the *sensu stricto* clade to which also *Lm* belongs. These include the pathogenic species *L. ivanovii*, and the non-pathogenic species *L. innocua*, *L. marthii*, *L. seeligeri* and *L. welshimeri* (Schardt et al., 2017). Like *Lm*, they all formed VBNC subpopulations in mineral water (**Fig. 3A–J**). *L. ivanovii* exhibited the greatest drop in culturability (3 log) after 7 days (**Fig. 3A**), which meant that >99% of viable *L. ivanovii* cells present at day 7 were in a VBNC state (**Fig. 3F**). *L. marthii* showed transition dynamics similar to *Lm*, with a slower decline in culturability after 7 days (**Fig. 3C**, compare to **Fig. 1B** and **Extended Data Fig. 1A–D**). Finally, *L. innocua*, *L. seeligeri* and *L. welshimeri* displayed intermediate profiles of culturability loss (**Fig. 3B, D, E**) and VBNC cell formation (**Fig. 3G, I, J**). The anti-*Lm* antibody also reacted with the CW of these species, so we used it to follow their CW status by fluorescence microscopy. As observed with *Lm*, the other *Listeria* species lost their CW while transitioning to a VBNC state (**Fig. 3K–O, U**). In parallel, the anti-CWD *Lm* antibody was also able to reveal the gradual exposure of the plasma membrane in these species as they shed their CW (**Fig. 3P–U**).

**Fig. 3.**
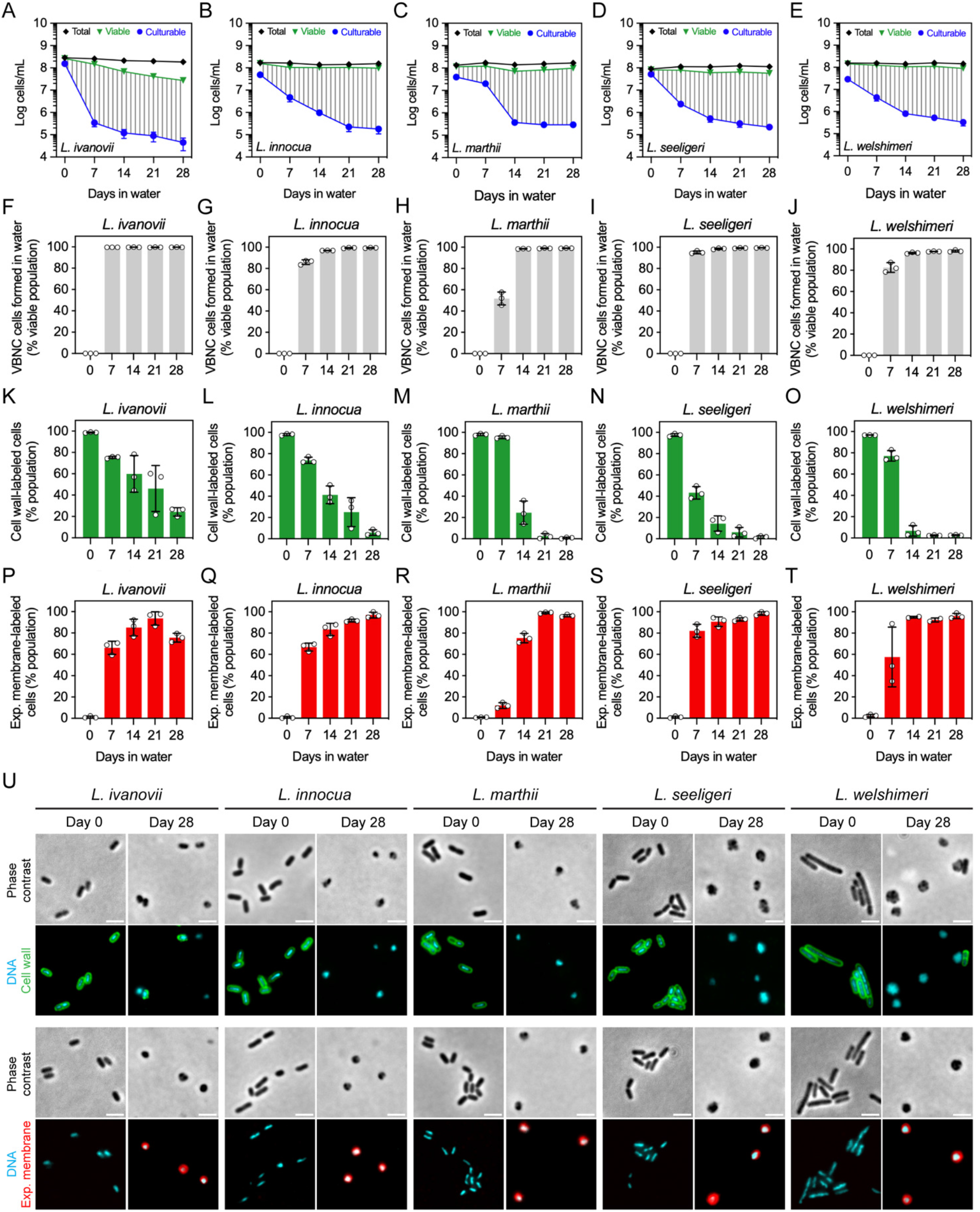
A CWD VBNC state occurs in other *Listeria* species. **(A–E)** Total, viable and culturable cell profiles of mineral water suspensions (10^8^ cells/mL) of *Listeria sensu stricto* clade members *L. ivanovii* (A), *L. innocua* (B), *L. marthii* (C), *L. seeligeri* (D) and *L. welshimeri* (E). Culturable bacteria were determined by enumeration of CFUs, while total and viable bacteria were quantified by flow cytometry. Data are represented as the mean ± SD from three independent suspensions. The area covered with vertical grey dashes indicates the VBNC cell population. **(F–J)** Fraction of the *L. ivanovii* (F), *L. innocua* (G), *L. marthii* (H), *L. seeligeri* (I) and *L. welshimeri* (J) populations consisting of VBNC cells formed in mineral water. Data are represented as the mean ± SD from three independent suspensions, each indicated by a dot. **(K–T)** Fraction of the *L. ivanovii* (K, P), *L. innocua* (L, Q), *L. marthii* (M, R), *L. seeligeri* (N, S) and *L. welshimeri* (O, T) populations showing CW (K–O) or exposed plasma membrane (P–T) labelling (anti-*Lm* and anti-CWD *Lm*, respectively) by fluorescence microscopy at the indicated timepoints. Data are represented as the mean ± SD from three independent suspensions, each indicated by a dot. **(U)** Phase-contrast and fluorescence micrographs of the five *Listeria* species sampled from mineral water suspensions at the indicated timepoints. Bacteria were fixed and fluorescently labelled for visualization of DNA (Hoechst 33342), CW (anti-*Lm*) and exposed membrane (anti-CWD *Lm*). Scale bar: 2 µm.

Collectively, these results show that the emergence of CWD VBNC forms in mineral water is a transversal phenomenon in *Listeria sensu stricto* species.

### *Lm* changes the physicochemical properties of its plasma membrane to adapt to a CWD lifestyle in water

The bacterial CW protects shape and counters the intracellular osmotic pressure, protecting the cell from osmotic lysis. We sought to understand how CWD *Lm* cells can survive in a hypotonic medium, like mineral water, without signs of lysis. We hypothesized that *Lm* may change the properties of its plasma membrane to become more resistant to osmotic pressure before shedding its CW. Indeed, bacteria modulate the fluidity of their plasma membrane, in response to changing environmental factors, to preserve the physical and functional integrity of their interface with the external environment. This is mainly accomplished by changing the fatty acid (FA) composition of membrane phospholipids, which adjusts their degree of packing and, consequently, the fluidity of the membrane (Yoon et al., 2015).

We thus analyzed the FA composition of the *Lm* membrane by gas chromatography coupled to mass spectrometry. Between day 0 and day 28, we observed a decrease in the relative abundance of anteiso branched-chain species (a-BFA) a-C15:0 and a-C17:0 (**Extended Data Fig. 1**), which comprise the majority of the *Lm* FA population and are key regulators of membrane fluidity in Gram-positive bacteria (Yoon et al., 2015). In contrast, linear saturated (SFA) and unsaturated (UFA) FAs showed increased relative levels, mainly on account of C16:0 and C16:1 species (**Extended Data Fig. 1**). Due to their minor representation in the initial FA population, the fold increase in the SFA and UFA levels was substantial, when compared to the fold change of the more abundant a-BFAs (**Fig. 4A**). We then investigated if these changes in FA composition were associated with an alteration in the *Lm* membrane fluidity by measuring the generalized polarization (GP) of laurdan, a ratiometric probe that shifts its fluorescence emission peak in response to local membrane phase transitions caused by fluidity changes (Scheinpflug et al., 2017). An increase of the laurdan GP in labeled *Lm* cells during the first 14 days was suggestive of decreased membrane fluidity (**Fig. 4B**). To confirm this observation, we directly measured the fluidity-dependent diffusion of the fluorogenic dye Nile red in the *Lm* membrane using total internal reflection fluorescence correlation spectroscopy (TIR-FCS). This technique has been recently implemented to quantify bacterial membrane fluidity in Gram-positive bacteria (Barbotin et al., 2023). TIR-FCS showed a significant reduction of the diffusion coefficient of Nile red in rod-shaped cells between 7 and 14 days (**Fig. 4C**), suggestive of membrane rigidification. This corresponds to the period when *Lm* is most severely impacted by CW damage and loss (**Fig. 1K**). Notably, the diffusion coefficient in CWD coccoid cells was similar after 7 or 14 days of incubation in water (**Fig. 4C**), consistent with increased membrane packing to adapt to a wall-less lifestyle. Furthermore, after 14 days in water, the diffusion coefficient was similar in rod-shaped and coccoid cells, suggesting that the reduction in membrane fluidity occurs prior to CW loss.

**Fig. 4.**
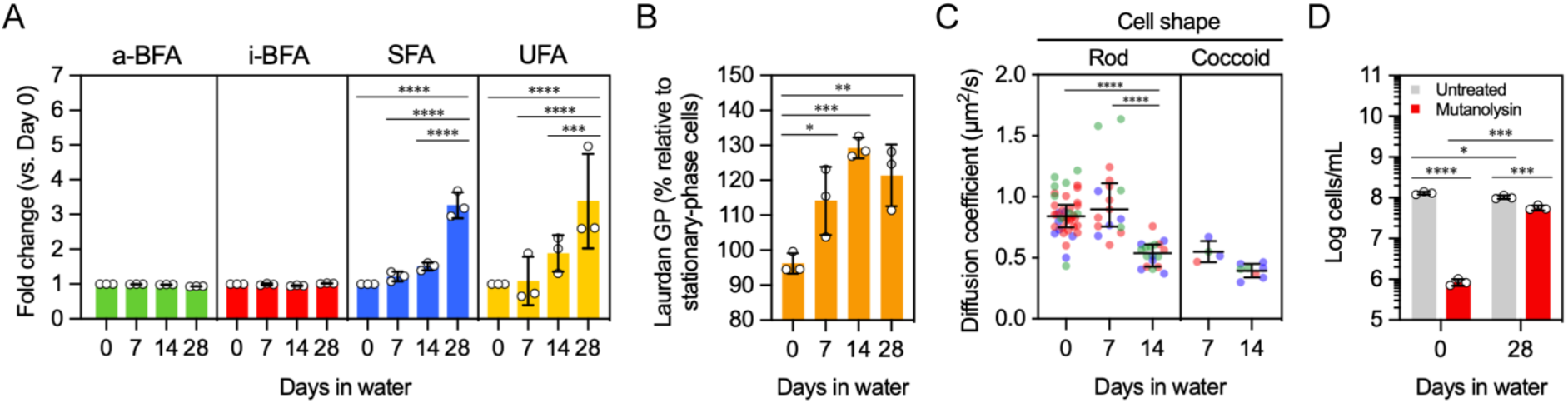
*Lm* adjusts its plasma membrane properties to adapt to a CWD lifestyle. **(A)** Fold change of the relative abundance of anteiso-branched (a-BFA), iso-branched (i-BFA), unsaturated (UFA) and saturated (SFA) fatty acids present in *Lm* EGDe sampled from mineral water suspensions (10^8^ cells/mL) at the indicated timepoints relative to the first timepoint. Fold change values were calculated from the data presented in Extended Data Fig. 1, and are represented as the mean ± SD from three independent suspensions. Statistical analysis was performed using a two-way ANOVA test with Tukey’s post hoc multiple comparison correction. ***, p≤0.001; ****, p≤0.0001. **(B)** Generalized polarization (GP) of laurdan-labeled *Lm* EGDe sampled from mineral water suspensions (10^8^ cells/mL) at the indicated timepoints. GP values were measured by fluorescence spectroscopy and normalized to those from overnight-grown bacteria. Increasing relative GP suggests a reduction of the bacterial membrane fluidity. Data are represented as the mean ± SD from three independent suspensions, each indicated by a dot. Statistical analysis was performed using a one-way ANOVA test with Dunnett’s post hoc multiple comparison correction. *, p≤0.05; **, p≤0.01; ***, p≤0.001. **(C)** Membrane diffusivity of Nile red in *Lm* EGDe cells sampled from mineral water suspensions at the indicated timepoints. Diffusion coefficient values were measured in Nile red-stained cells of rod and coccoid shape by total internal reflection fluorescence correlation spectroscopy (TIR-FCS). Data are represented as scattered dot plots with the median ± interquartile range from three independent suspensions. Each dot represents one measured bacterium and bacteria from the same suspension are indicated by the same color. Statistical analysis was performed using a one-way ANOVA test with Tukey’s post hoc multiple comparison correction. ****, p≤0.0001. **(D)** Osmotic sensitivity of CWD *Lm* EGDe adapted versus non-adapted to mineral water. The total bacterial population in mineral water suspensions treated or not with mutanolysin at the indicated timepoints was quantified by flow cytometry. Data are represented as the mean ± SD from three independent suspensions. Statistical analysis was performed using a two-way ANOVA test with Šidák’s post hoc multiple comparison correction. *, p≤0.05; ***, p≤0.001; ****, p≤0.0001.

Altogether, these findings indicate that *Lm* alters the physicochemical properties of its plasma membrane while transiting to a CWD VBNC state in mineral water. As a result of changes in FA composition, although not excluding the contribution of other membrane components, the membrane becomes more rigid, which may protect the wall-less bacterial cell from osmotic lysis. In agreement with this hypothesis, total *Lm* numbers were notably reduced (2 log) when freshly prepared suspensions were immediately treated with mutanolysin, which digests the *Listeria* CW, without allowing the *Lm* cells to adapt to the hypotonic medium (**Fig. 4D**). In contrast, bacteria from 28-day-old suspensions were insensitive to this treatment (**Fig. 4D**).

### Transcriptomics highlight the role of stress response in the formation of CWD VBNC *Lm*

Despite several studies reporting the induction of a VBNC state in *Lm* under different stressful conditions (Besnard et al., 2002; Bremer et al., 1998; Cunningham et al., 2009; Highmore et al., 2018; Lindbäck et al., 2010; Noll et al., 2020; Robben et al., 2018), the molecular factors and pathways involved in this transition remain elusive.

To identify early effectors required for VBNC state transition in mineral water, we analyzed the transcriptional changes in *Lm* cells after 7 days, when loss of culturability and CW alterations are first observed (**Fig. 1**). RNA-seq analysis identified a total of 1229 differentially expressed genes (q-value ≤ 0.05, absolute log_2_ fold change ≥ 1), of which 593 were downregulated and 636 were upregulated (**Fig. 5A; Supplementary Table 2**). Gene set enrichment analysis revealed the most prevalent upregulated and downregulated biological processes and pathways. Downregulated genes were found associated with biosynthesis of nucleotides and coenzymes (biotin, pyridoxal phosphate, coenzyme A), transcription regulation, uptake of phosphate and carbohydrates (maltose/maltodextrin and trehalose phosphotransferase systems), cell envelope assembly (biosynthesis of glycerophospholipids and teichoic acids) and maintenance (peptidoglycan catabolism), cell division (division septum assembly), energy production (pyruvate metabolism, ATP synthesis-coupled proton transport), and protein secretion (**Fig. 5B**, **Extended Data Fig. 1A–C**). Upregulated genes were linked with acquisition and/or metabolism of amino acids (aspartate, glutamate, methionine, cysteine, isoleucine, valine, leucine, threonine, arginine); biosynthesis of pyrimidine nucleotides, uptake of carbohydrates (glucose/mannose phosphotransferase systems) and metal ions (iron, zinc); protein translation and folding, and response to osmotic (transport of compatible solutes carnitine/glycine betaine) and oxidative stress (glutathione metabolism) (**Fig. 5C; Extended Data Fig. 1D–F**). These results are consistent with a physiological transition taking place in a population of mixed culturable states. Most downregulated genes likely reflect the transition from a vegetative growth state to a VBNC state, whereas most upregulated genes possibly mirror bacterial responses to nutritional and hypoosmotic stresses.

**Fig. 5.**
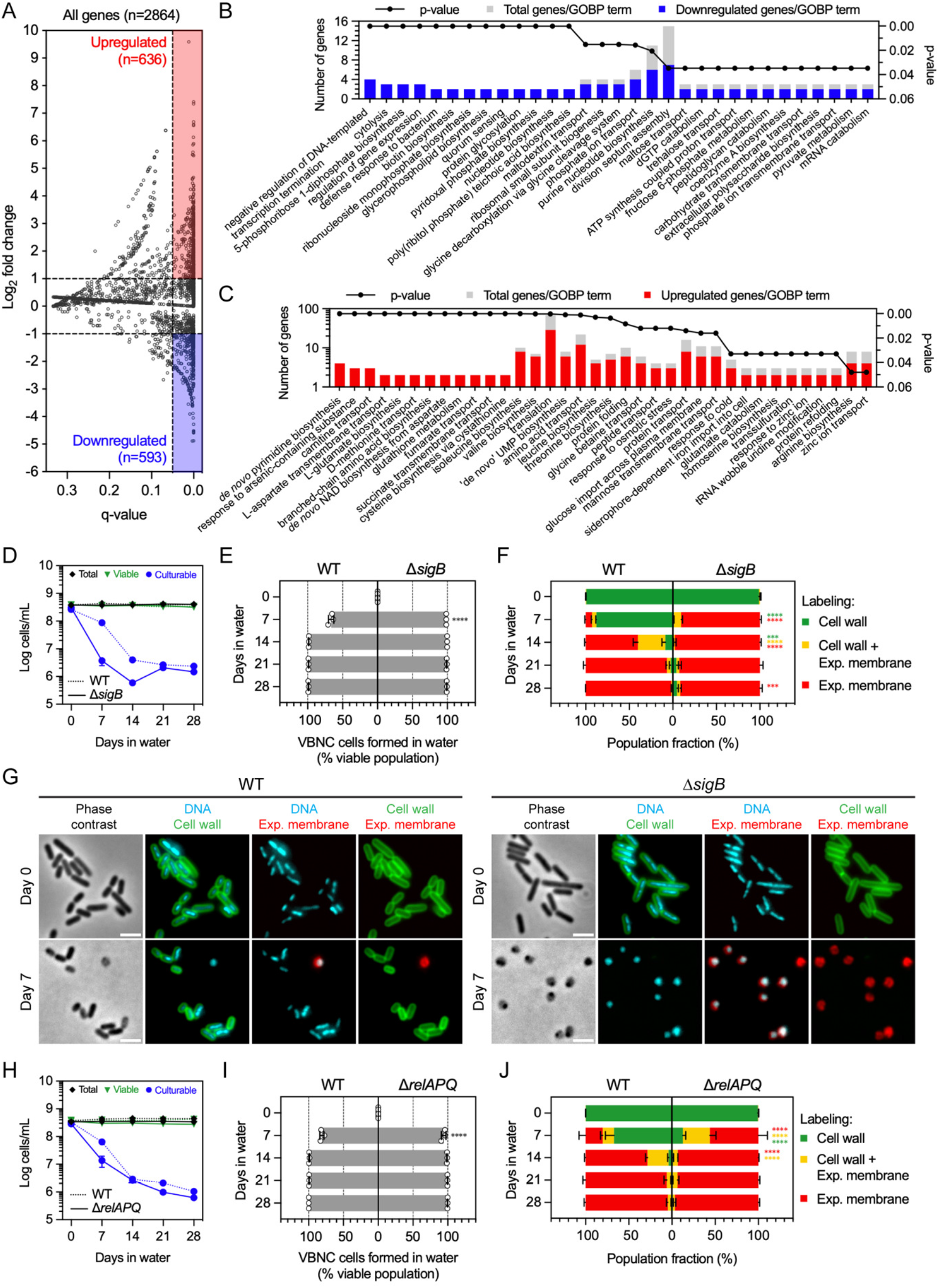
Transcriptional profiling reveals a major regulatory role for SigB and stress response mechanisms in the onset of VBNC *Lm* formation. **(A)** Scatter plot of *Lm* EGDe protein-coding genes (n=2864) according to the statistical significance (q-value) and magnitude of transcriptional change (log2 fold change) determined at day 7 relative to day 0 by RNA-seq. Horizontal dashed lines indicate fold change thresholds (|log_2_|=1) and vertical dashed line indicates statistical significance threshold (q-value=0.05). Genes statistically considered to be differentially regulated (n=1229) are grouped into 636 upregulated genes (red box) and 593 downregulated genes (blue box). **(B, C)** Functional analysis of statistically enriched gene ontology biological process (GOBP) terms in downregulated (B) and upregulated genes (C). GOBP terms are ranked from left to right by increasing p-value and decreasing fraction of differentially regulated genes per term. **(D)** Total, viable and culturable cell profiles of mineral water suspensions of wild type (WT) and SigB-deficient (Δ*sigB*) *Lm* EGDe. Culturable bacteria were determined by enumeration of CFUs, while total and viable bacteria were quantified by flow cytometry. Data are represented as the mean ± SD from three independent suspensions. **(E)** Fraction of the WT and Δ*sigB* populations consisting of VBNC cells formed in mineral water. Data are represented as the mean ± SD from three independent suspensions, each indicated by a dot. Statistical analysis between mutant and WT strains was performed using a two-way ANOVA test with Šidák’s post hoc multiple comparison correction. ****, p≤0.0001. **(F)** Fraction of the WT and Δ*sigB* populations showing single- or double-labeling of CW (WGA) and exposed plasma membrane (anti-CWD *Lm*) by fluorescence microscopy, at the indicated timepoints. Data are represented as stacked bars (one for each labelling group) with mean ± SD from three independent suspensions. Statistical analysis between Δ*sigB* and WT strains was performed for each labelling group using a two-way ANOVA test with Tukey’s post hoc multiple comparison correction. ***, p≤0.001; ****, p≤0.0001. **(G)** Phase-contrast and fluorescence micrographs of WT and Δ*sigB* bacteria sampled from mineral water suspensions at the indicated timepoints. Bacteria were fixed and fluorescently labelled for visualization of DNA (Hoechst 33342), CW (WGA) and exposed plasma membrane (anti-CWD *Lm*). Scale bar: 2 µm. **(H)** Total, viable and culturable cell profiles of mineral water suspensions of wild type (WT) and RelAPQ-deficient (Δ*relAPQ*) *Lm* EGDe. Culturable bacteria were determined by enumeration of CFUs, while total and viable bacteria were quantified by flow cytometry. Data are represented as the mean ± SD from three independent suspensions. **(I)** Fraction of the WT and Δ*relAPQ* populations consisting of VBNC cells formed in mineral water. Data are represented as the mean ± SD from three independent suspensions, each indicated by a dot. Statistical analysis between mutant and WT strains was performed using a two-way ANOVA test with Šidák’s post hoc multiple comparison correction. ****, p≤0.0001. **(J)** Fraction of the WT and Δ*relAPQ* populations showing single- or double-labeling of CW (WGA) and exposed plasma membrane (anti-CWD *Lm*) by fluorescence microscopy, at the indicated timepoints. Data are represented as stacked bars (one for each labelling group) with mean ± SD from three independent suspensions. Statistical analysis between Δ*relAPQ* and WT strains was performed for each labelling group using a two-way ANOVA test with Tukey’s post hoc multiple comparison correction. ****, p≤0.0001.

Interestingly, prophage loci were almost completely activated and among the most strongly upregulated genes (**Supplementary Table 3**), in line with previous reports linking prophage activation with environmental stress (Argov et al., 2019; Duru et al., 2021; Ivy et al., 2012; Wang et al., 2010). In agreement with a stress response activation, nearly half of the regulon controlled by the stress-responsive sigma factor SigB (181 out of 455 genes) was induced (**Supplementary Table 4**). This elevated number of upregulated SigB-controlled genes prompted us to investigate its involvement in the transition of *Lm* to a VBNC state. We found that SigB-deficient *Lm* cells transitioned considerably faster than wild-type cells, with a 2-log decline in CFU counts resulting in >90% of viable cells in a non-culturable state after 7 days (**Fig. 5D, E**). Importantly, >90% of the *Lm* population had already converted to CWD coccoid forms (**Fig. 5F, G**). These results reveal a major modulating role for SigB in *Lm* adaptation to nutritional deprivation and generation of CWD VBNC forms in mineral water. The unaffected viability of Δ*sigB* cells indicates, however, that SigB is not essential for *Lm* survival in this situation (**Fig. 5D**).

The stringent response is an important stress signaling mechanism that regulates adaptation to starvation via the alarmone (p)ppGpp (Irving et al., 2021). The production of (p)ppGpp was shown to be promoted during transition to a VBNC state, and (p)ppGpp-deficient bacteria were found to lose culturability at a higher rate than their wild-type counterparts (Bai et al., 2021; Boaretti et al., 2003). Although our transcriptomic data showed no upregulation of the alarmone synthase-encoding genes (*relA*, *relP* and *relQ*) (**Supplementary Table 2**), we examined if the enzymatic activity of these proteins could impact the transition to a VBNC state. Compared to wild type *Lm*, a Δ*relAPQ* strain displayed a significantly faster transition after 7 days (**Fig. 5H, I**), which was correlated with a faster decline of the walled population in the same period (**Fig. 5J**). This result suggest that the stringent response plays a role in the early phase of VBNC *Lm* formation.

### The autolysin NamA is a major mediator of *Lm* CW loss and VBNC state entry

To gain a molecular insight into the *Lm* CW remodeling dynamics involved in VBNC *Lm* formation in water, we then focused on genes involved in CW metabolism. These genes presented a heterogeneous expression profile, showing either down/upregulation or no change (**Supplementary Table 5**). Peptidoglycan maturation and turnover is carried out by a family of peptidoglycan hydrolases, commonly called autolysins, that cleave different bonds within the peptidoglycan structure (Höltje, 1998). *Lm* encodes around 20 proteins with confirmed or predicted autolytic activity (Bierne & Cossart, 2007; Popowska & Markiewicz, 2006). Our transcriptomic data indicated that many known and putative *Lm* autolysin-coding genes were strongly downregulated after 7 days (e.g. *lmo0394, p60, aut, lmo1215, lmo1521, lmo2522, ami, namA*) (**Supplementary Table 5**), suggesting that degradation of the CW prior to shedding is carried out efficiently by an existing autolytic activity, without need for additional protein synthesis. We thus tested *Lm* mutants of genes encoding autolysins with different classes of bond-cleaving activity: the DL-endopeptidase p60/Iap, the *N*-acetylmuramoyl-L-alanine amidase Ami, the *N*-acetylglucosaminidases Auto and NamA, and the putative *N*-acetylmuramidases/lytic transglycosylases and resuscitation-promoting factor (Rpf)-like proteins Lmo0186 and Lmo2522 (Bierne & Cossart, 2007; Carroll et al., 2003; Pinto et al., 2013).

*Lm* deficient in p60 (**Extended Data Fig. 1A–C**) or in both Rpf proteins (**Extended Data Fig. 1D–F**) showed culturability, VBNC cell formation and CW loss profiles largely similar to those of wild type bacteria. Ami-deficient *Lm* showed a higher proportion of walled bacteria at day 14, associated with a delay in culturability decline and formation of VBNC cells at day 7 (**Extended Data Fig. 1G–I**). Interestingly, in the absence of Auto, *Lm* presented a significant drop in culturable and walled cell numbers – and thus larger VBNC population – at day 7 (**Extended Data Fig. 1J–L**).

The most striking phenotype was observed with NamA-deficient *Lm*, which displayed strongly delayed dynamics of transition to a VBNC state (**Fig. 6A–C**). Indeed, the culturability of Δ*namA* bacteria was barely affected at day 7 and showed 10-fold higher values than wild type bacteria at day 14 (**Fig. 6A**). Importantly, more than 90% of the Δ*namA* population still conserved their CW after 14 days, compared to 44% of wild type *Lm* (**Fig. 6C**). As the export of NamA to the bacterial surface is specifically mediated by the accessory Sec system ATPase SecA2 (Lenz et al., 2003), we investigated whether SecA2-deficient *Lm* exhibited a phenotype similar to NamA-deficient *Lm*. Indeed, Δ*namA* and Δ*secA2* bacteria demonstrated very similar dynamics with respect to culturability loss, VBNC cell formation and CW loss, despite substantially lower initial culturable Δ*secA2* counts (**Fig. 6D–F**). This initial difference is likely due to a division/scission defect of the Δ*secA2* strain that results in a chaining phenotype (**Fig. 6G**), as SecA2 mediates the secretion of both NamA and p60 autolysins (Machata et al., 2005). However, the lack of a role for p60 in CW shedding during VBNC *Lm* formation (**Extended Data Fig. 1A–C**) suggests that the Δ*secA2* phenotype is caused by the absence of exported NamA. Supporting this hypothesis, CW loss by Δ*secA2* cells was delayed to the same extent as in Δ*namA* cells (**Fig. 6C, F**).

**Fig. 6.**
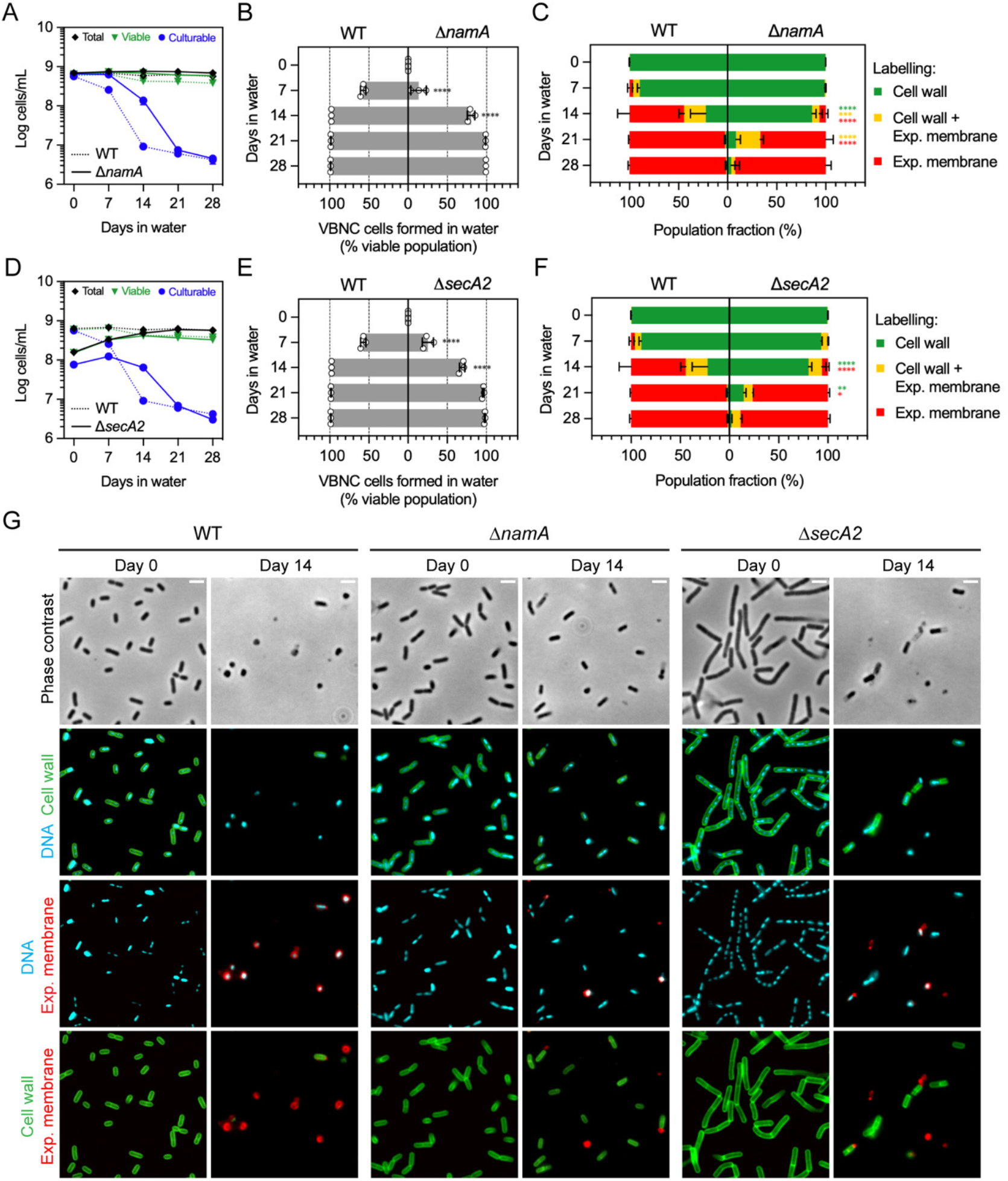
The autolysin NamA promotes *Lm* CW shedding and VBNC entry. **(A, D)** Total, viable and culturable cell profiles of mineral water suspensions of wild type (WT) and NamA-deficient (A) or SecA2-deficient (D) *Lm* EGDe. Culturable bacteria were determined by enumeration of CFUs, while total and viable bacteria were quantified by flow cytometry. Data are represented as the mean ± SD from three independent suspensions. **(B, E)** Fraction of the WT and Δ*namA* (B) or Δ*secA2* (E) populations consisting of VBNC cells formed in mineral water. Data are represented as the mean ± SD from three independent suspensions, each indicated by a dot. Statistical analysis between mutant and WT strains was performed using a two-way ANOVA test with Šidák’s post hoc multiple comparison correction. ****, p≤0.0001. **(C, F)** Fraction of the WT and Δ*namA* (C) or Δ*secA2* (F) populations showing single- or double-labeling of CW (WGA) and exposed plasma membrane (anti-*Lm*_CWD_) by fluorescence microscopy, at the indicated timepoints. Data are represented as stacked bars (one for each labelling group) with mean ± SD from three independent suspensions. Statistical analysis between mutant and WT strains was performed for each labelling group using a two-way ANOVA test with Tukey’s post hoc multiple comparison correction. *, p≤0.05; **, p≤0.01; ***, p≤0.001; ****, p≤0.0001. **(G)** Phase-contrast and fluorescence micrographs of WT, Δ*namA* and Δ*secA2 Lm* sampled from mineral water suspensions (10^8^ cells/mL) at the indicated timepoints. Bacteria were fixed and fluorescently labelled for visualization of DNA (Hoechst 33342), CW (WGA) and exposed plasma membrane (anti-CWD *Lm*). Scale bar: 2 µm.

Altogether, these results reveal the *Lm* autolysin NamA as a major player in VBNC *Lm* formation, with an important role in the events tied to CW breakdown and shedding.

### CWD VBNC *Lm* can revert back to a vegetative, walled and virulent state

Some bacteria, mostly Gram-negative species, have been shown to exit the VBNC state and regain culturability under specific “resuscitation” conditions (Ayrapetyan et al., 2018; Dong et al., 2020). Reversion of the VBNC state in *Lm* and other Gram-positive bacteria remains a challenge (Dong et al., 2020; Lotoux et al., 2022). Revival attempts through nutrient supplementation (e.g. inoculation into fresh or conditioned medium, pure or diluted) were unsuccessful in our hands (our unpublished results). We thus turned to the chicken embryo model, which was previously used to effectively revive VBNC bacteria, including *Lm* (J. M. Cappelier et al., 1999; Jean Michel Cappelier et al., 2007; Chaveerach et al., 2003; Talibart et al., 2000). We inoculated embryonated chicken eggs with a suspension of GFP-expressing VBNC *Lm* containing 10^6^ viable cells/mL, of which <1 cells/mL were culturable. In parallel, eggs inoculated with mineral water or with vegetative *Lm* from an overnight broth culture served respectively as negative and positive controls. Two days after inoculation, eggs were processed to assess the presence of culturable *Lm*. All embryonated eggs inoculated with VBNC *Lm* scored positive for bacterial growth (24 out of 24 eggs), similarly to embryonated eggs inoculated with vegetative *Lm* (8 out of 8 eggs). As expected, no bacterial growth was observed in eggs inoculated with mineral water (**Supplementary Table 6**).

A critical point with resuscitation of VBNC bacteria is whether the recovered culturable cells resulted from a true revival of VBNC forms or from regrowth of a trace number of culturable cells. To rule out the latter possibility, VBNC *Lm* were inoculated in parallel in BHI, a rich medium that does not support VBNC *Lm* resuscitation (Besnard et al., 2002). This resulted in bacterial growth in only 9.52% (8 in 84) of inoculated wells, which largely contrasted with growth obtained from 100% of inoculated embryonated eggs. This significantly different proportion of *Lm* growth before and after passage of VBNC cells in embryonated eggs (p=1.63×10^−17^) attests the successful resuscitation of *Lm* from the VBNC state (**Supplementary Table 6**). As a further control, VBNC cells were also inoculated into non-embryonated eggs, which were shown to fail in promoting VBNC *Lm* revival (Jean Michel Cappelier et al., 2007). Whereas vegetative *Lm* were able to proliferate in non-embryonated eggs, we observed no growth coming from VBNC *Lm*-inoculated eggs (**Supplementary Table 6**), underlining the requirement of an embryo for VBNC *Lm* resuscitation and further supporting that the revival of VBNC *Lm* in embryonated eggs was not due to residual culturable bacteria in the inoculum.

To confirm if these awakened *Lm* were phenotypically equivalent to vegetative *Lm*, we investigated their morphology and virulence. Fluorescence microscopy of two revived *Lm* clones revealed populations consisting of GFP-expressing, walled, rod-shaped cells that are indistinguishable from vegetative bacteria grown in broth medium (**Fig. 7A**). We then assessed the virulence of the revived clones by infecting human trophoblastic (JEG-3) and hepatocytic (HepG2) cell lines. Quantification of the intracellular bacterial load over time revealed no differences between vegetative and revived *Lm* (**Fig. 7B**). Microscopy analysis of infected cells showed that both revived *Lm* clones produced foci of infected cells after 6 h and spread to the rest of the cell monolayer by 72 h post-infection as efficiently as vegetative *Lm* (**Fig. 7C**), supported by an equal capacity of polymerizing host actin into propulsive comet-like tails (**Fig. 7D**).

**Fig. 7.**
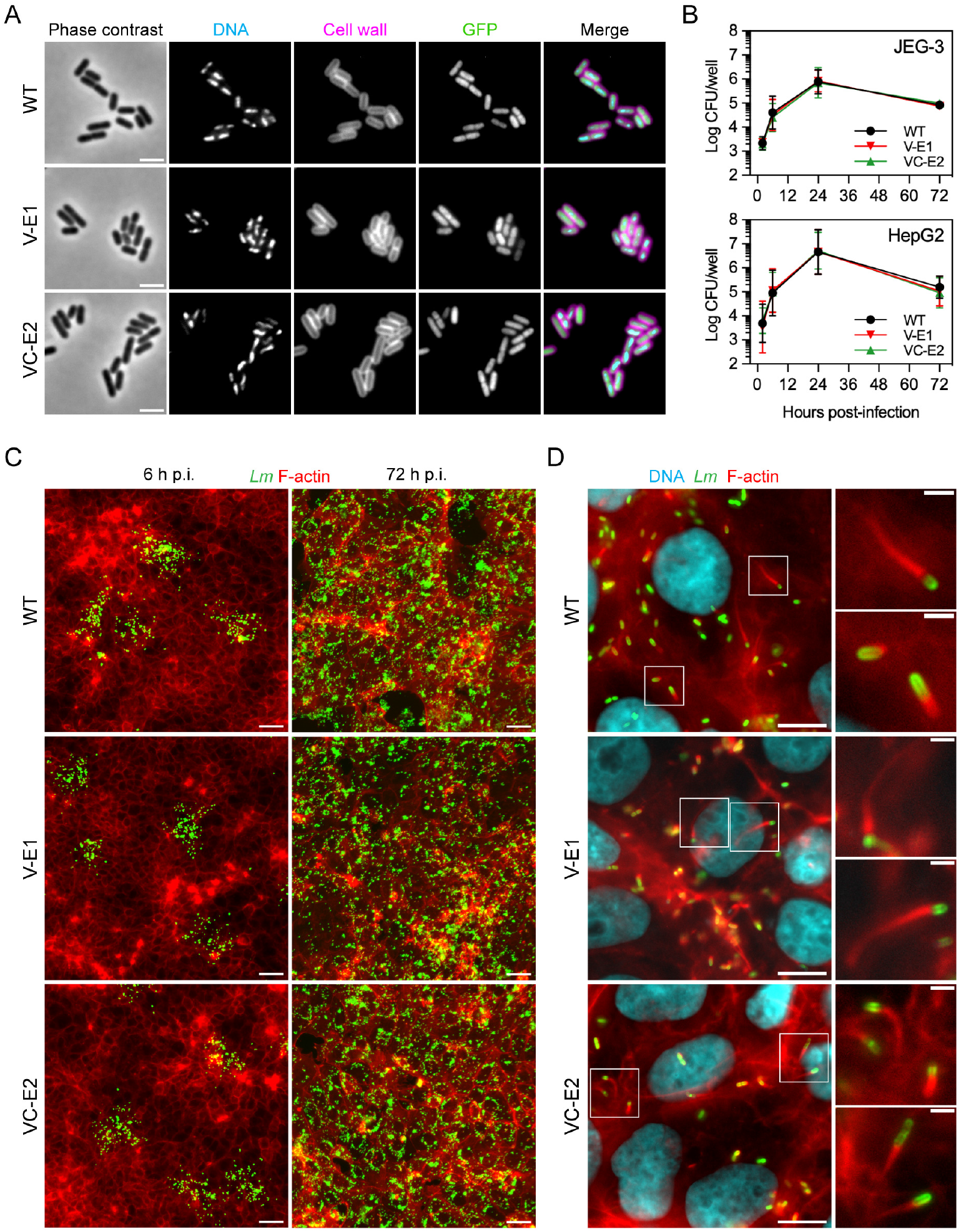
Chicken embryo passage restores culturability and virulence to VBNC *Lm*. **(A)** Phase-contrast and fluorescence micrographs of GFP-expressing *Lm* EGDe before (WT) and after incubation in mineral water and passage through embryonated chicken eggs (clones V-E1 and VC-E2). Bacteria were fixed and fluorescently labelled for visualization of DNA (Hoechst 33342) and CW (WGA). Scale bar: 2 µm. **(B)** Intracellular replication of WT and V-E1 and VC-E2 *Lm* in human epithelial JEG-3 and HepG2 cell lines. At each timepoint, intracellular bacteria were determined by enumeration of CFUs coming from the plating of epithelial cell lysates. Data are represented as mean ± SD from two independent infection assays. **(C)** Low-magnification fluorescence micrographs of JEG-3 cell monolayers infected with WT, VE-1 and VC-E2 strains showing the cell-to-cell spread of intracellular bacteria between 6h (isolated clusters of infected cells) and 72h post-infection (generalized infection of the cell monolayer). Cells were fixed and fluorescently labelled for visualization of bacteria (anti-*Lm*) and F-actin (phalloidin). Bacteria-derived fluorescence signal was digitally enhanced for clarity purposes. Scale bar: 50 µm. **(D)** Fluorescence micrographs of WT, VE-1 and VC-E2 *Lm* in the cytoplasm of infected JEG-3 cells at 6h post-infection. Cytosolic bacteria polymerize host actin to form comet-like tail structures that promote bacterial motility inside cells and subsequent intercellular spread. Infected cells were fixed and fluorescently labelled for visualization of DNA (Hoechst 33342), bacteria (anti-*Lm*) and F-actin (phalloidin). Bacteria highlighted by white inset squares are shown enlarged in right-side panels (nuclei are not shown for better visualization of bacteria and actin tails). Scale bars: 10 µm (left-side panels), 2 µm (right-side panels).

Altogether, these results provide strong evidence that the CWD VBNC state of *Lm* is fully reversible and bacteria are able to switch back to a walled and cell-infecting state.

## Discussion

*Lm* is a ubiquitous bacterium known to be tolerant to several biotic and abiotic insults. A key factor explaining the presence and survival of this non-sporulating species in a multiplicity of harmful environments is the ability to phase into a dormant VBNC state. First investigated over 20 years ago (Besnard et al., 2000a), the VBNC state of *Lm* remains however largely uncharacterized. In this work, we showed that *Lm* switches from rod-shaped to coccoid cell as it transitions to a VBNC state in mineral water. We further revealed that this coccoid cell form represents a CWD variant that is generated by a molting-like process of CW shedding. Our findings suggest that generation of CWD cells is key for the transition of *Lm* into a non-culturable dormant state in a natural water environment.

The major finding of this work is that CWD VBNC bacteria can naturally emerge and persist in a hypotonic (i.e. osmotically hostile) environment. Following an adaptation period of the initial walled cells to mineral water, the CWD *Lm* forms described in this work are relatively robust in this environment without osmoprotection. This contrasts with other CWD bacterial types, such as L-forms or the recently reported actinomycete S-cells (Ramijan et al., 2018), which are formed in an osmoprotective (i.e. hypertonic) environment, or require one during their formation, to avoid explosive cell lysis after CW loss (Claessen & Errington, 2019). We also showed that, as *Lm* transitions to a CWD VBNC state, it fine-tunes the physicochemical characteristics of its plasma membrane to become more rigid and potentially more resistant to lysis. It is also possible that physical changes in the cytoplasm could account for the increased mechanical resistance to the extracellular hypoosmotic conditions. To this regard, low metabolic activity has been shown to induce a “glassy” behavior of the cytoplasm that might help bacteria to preserve their cellular architecture (Parry et al., 2014).

The CWD *Listeria* described in this study are also distinct from L-forms in their mechanism of formation. Whereas *in vitro*-generated L-forms are typically induced by artificial weakening/breakdown of the CW (e.g. exposure to antibiotics or lytic enzymes), we show that CWD VBNC *Listeria* arise naturally in mineral water. S-cells were also shown to be naturally formed in response to hyperosmotic conditions; however, unlike CWD VBNC *Listeria*, they did not survive in a hypotonic medium (Ramijan et al., 2018). Notably, *Lm* is ubiquitously found in oligotrophic aquatic environments and its occurrence in environmental surface water samples, for example, is estimated at 10–30% after culturing in selective rich medium (Lyautey et al., 2007; Raschle et al., 2021; Sharma et al., 2020; Weller et al., 2015). The presence of unculturable CWD *Lm* in these environmental niches may therefore be vastly underestimated. In this regard, the antibodies generated in this study that specifically recognize wall-less VBNC forms of different *Listeria* species, including the two pathogenic *Lm* and *L. ivanovii*, constitute a novel biomolecular tool that can be potentially used in the detection of dormant pathogens, otherwise untraceable by standard growth-based techniques.

Cryo-ET characterization of the *Lm* CW shedding process showed the extrusion of the bacterial protoplast through one of many breaches in the CW sacculus. A similar process was notably first observed in *Bacillus subtilis* cells transitioning to an L-form state (Domínguez-Cuevas et al., 2012), which suggests common mechanisms underlying the CW loss event in both species. Indeed, perturbations in CW loss affected the production of *B. subtilis* L-forms (Domínguez-Cuevas et al., 2012) as well as of VBNC *Lm* in water. Mutations in *B. subtilis* genes resulting in sustained autolysin activity and/or septum malformation were found to promote CW extrusion and L-form emergence (Domínguez-Cuevas et al., 2012). Interestingly, we identified the *Lm* autolysin NamA as an important player in this process, since NamA-deficient cells were strongly delayed in CW shedding and VBNC state entry. Among the other tested autolysins, only Ami promoted the *Lm* CW shedding process, although at a less significant level compared to NamA. An in-depth screening of the *Lm* autolysin collection should reveal the full list of peptidoglycan-degrading enzymes involved in the formation of CWD VBNC *Lm*.

Inactivation of the *Lm* general stress response or the stringent response, via genetic deletion of the transcription factor SigB or the (p)ppGpp synthetases (RelAPQ), resulted in unexpectedly faster dynamics of VBNC cell formation. This phenotype was particularly strong in SigB-deficient cells, almost completely wall-less after just 7 days. Intriguingly, the absence/misregulation of these important stress response systems did not affect *Lm* cell viability, indicating that they are not required for bacterial survival in a rather stressful context. These results suggest that stress response regulators may secure a balance between vegetative and dormant states; in mineral water, the absence of these regulators might break the balance in favor of transition to the VBNC state. It will be interesting to understand whether the lack of SigB or a functional stringent response in these mutant bacteria has deleterious consequences, for example in their capacity to successfully exit the VBNC state.

We then showed that CW shedding during transition to a VBNC state is not limited to *Lm* and can occur in other pathogenic and non-pathogenic *Listeria* species. This may thus represent an evolutionary strategy within the *Listeria* genus to withstand prolonged nutritional deficiency. It has been hypothesized that the bacterial CW primarily evolved as a structure for storage of sugar- and amino acid-rich components (Claessen & Errington, 2019). Breakdown and salvage of CW components could constitute a bacterial mechanism to secure nutrients to sustain a minimal metabolic flux through the unknown duration of a VBNC state. Whether this ability to form wall-less dormant forms in similar conditions also extends to other species from phylogenetically related genera, including sporulating species, remains to be investigated.

Many pathogenic bacteria have been reported to transition to a VBNC state and, for some, this is associated with a loss of virulence (Zhao et al., 2017). We showed that CWD VBNC *Lm* reverted to a culturable state and recovered its CW and virulence, after passing through the chicken embryo. This indicates that one or more yet-unidentified signals present in this host environment can “wake up” these dormant *Lm* forms. Similar resurrection signals might be present in other eukaryotic hosts and in nature, and their identification constitutes a challenging but exciting research avenue.

It is becoming clear that a CWD state is an alternative lifestyle that enables bacteria to survive under stress and even proliferate without the mechanical protection of a rigid CW (Claessen & Errington, 2019; Ramijan et al., 2018). The emergence of CWD forms in phylogenetically distant bacterial species (Dannenberg et al., 2022; Ramijan et al., 2018; Slavchev et al., 2013) raises the possibility that transient CW loss may be a more common phenomenon among bacteria than expected. In a world dominated by walled microbes, a temporary wall-less state of dormancy might represent a strategy to promote bacterial persistence under harsh environmental contexts.

## Methods

### Bacteria, cell lines, growth media and conditions

Bacteria used in this work are listed in Supplementary Table 7 and were grown at 37 °C in brain and heart infusion (BHI) broth (with agitation) and agar media (BD Difco). Cell lines of human origin used in this work included JEG-3 trophoblasts (ATCC HTB-36), cultivated in MEM (Gibco, Thermo Fisher Scientific) supplemented with 10% (v/v) fetal calf serum (Eurobio Scientific), and HepG2 hepatocytes (ATCC HB-8065), cultivated in DMEM (Gibco, Thermo Fisher Scientific) supplemented with 10% (v/v) fetal calf serum. Cells were incubated at 37 °C in a humidified (90–95%) atmosphere with CO_2_ at 5% (for cell propagation) or 10% (for infected cells).

### Preparation of bacterial suspensions in mineral water

Bacterial suspensions in mineral water were prepared with bacteria from overnight-grown stationary-phase cultures. For each tested species, the bacterial concentration of stationary-phase cultures was determined beforehand by enumeration of colony-forming units (CFU) after plating in agar media. Bacteria were pelleted by centrifugation (3,000 × *g*, 3 min) and washed with 1 volume of sterile-filtered (0.22 µm) mineral water (henceforth referred simply as “mineral water”) for three times before resuspension in 1 volume of mineral water. Washed bacteria were then set to the desired concentration in a final volume of 30 mL of mineral water, and incubated statically at room temperature in an upright-standing, sterile tissue-culture flask (25 cm^2^, vented cap). Samples were collected for downstream analyses immediately after preparation (“day 0”) and after 7, 14, 21 and 28 days.

### Bacterial culturability and viability assays

Bacterial suspensions were regularly sampled for monitorization of the total, viable and culturable cell populations. The culturable population was quantified through enumeration of CFU following the plating of serial dilutions of the suspension on agar media. The total and viable populations were quantified by flow cytometry using a CytoFLEX S analyzer (Beckman Coulter) equipped with three excitation lasers (405, 488 and 561 nm) and operated by the CytExpert software (Beckman Coulter).

Suspensions prepared at 10^8^ cells/mL were ten-fold diluted in mineral water before acquisition at a flow rate of 10 µL/min. Bacteria-associated events were detected in a forward scatter (FSC) versus side scatter (SSC) plot (**Extended Data Fig. 1A**) and the total population was quantified by enumeration of FSC/SSC-gated events in a defined sample volume (10 µL). For determination of viable population using viability dyes, diluted suspensions were incubated in the dark either with 5(6)-carboxyfluorescein diacetate (CFDA, 30 µM) (Sigma-Aldrich) for 30 min or with a mix of SYTO 9 (3.34 µM) and propidium iodide (PI, 20 µM) from the LIVE/DEAD BacLight Bacterial Viability kit (#L7012, Molecular Probes, Thermo Fisher Scientific) for 15 min. Fluorescence emission by CFDA, SYTO 9 (525/40 nm bandpass) and PI (690/50 nm bandpass) was detected from FSC/SSC-gated bacteria, and populations containing viable (i.e. CFDA^+^ or SYTO 9^+^/PI^−^) or injured/dead bacteria (i.e. CFDA^−^ or PI^+^) were gated with the help of a “dead bacteria” control sample consisting of heat-treated (95 °C, 30 min) bacterial suspension (**Extended Data Fig. 1A**). The viable population was quantified by enumeration of CFDA^+^ or SYTO 9^+^/PI^−^-gated events in a defined sample volume (10 µL).

### Intracellular ATP quantification

The intracellular ATP content of bacteria suspended in mineral water was determined using the luciferase-based BacTiter-Glo Microbial Cell Viability Assay kit (Promega). As per the manufacturer instructions, 100 µL of bacterial suspension were mixed in an opaque white 96-well plate with 100 µL of room temperature-equilibrated BacTiter-Glo Reagent, and incubated in the dark for at least 5 min. Relative luminescence units (RLU) were then recorded in an Infinite M200 microplate reader (Tecan) with a 1-second integration time per well. Wells containing mineral water were used to obtain background luminescence.

### Peptidoglycan extraction and UHPLC analysis

Peptidoglycan was extracted from *Lm* EGDe as described (Sun et al., 2021). Bacteria (10^11^ cells) were harvested from mineral water suspensions (10^8^ cells/mL) on the day of preparation (day 0) and after 7 and 28 days by centrifugation (4,000 × *g*, 5 min), flash-frozen in liquid nitrogen and stored at −80 °C until further processing. Each bacterial cell pellet was then resuspended in 40 mL of cold distilled water, boiled for 10 min, cooled, and centrifuged. After suspending the cell pellet in 1 mL of distilled water, 1 mL of SDS solution (10% SDS in 100 mM Tris-HCl pH 7.0) at 60 °C was added and the suspension was boiled for 30 min and centrifuged (20 min, 25,000 × *g*). The pellet was resuspended in 2 mL of lysis solution (4% SDS in 50 mM Tris-HCl pH 7.0), boiled for 15 min, and washed six times with 60 °C-heated distilled water. Next, the pellet was treated with 2 mg/mL of pronase from *Streptomyces griseus* (Roche) in 50 mM Tris-HCl pH 7.0 for 1.5 h at 60°C, and afterwards with 10 μg/mL of DNase (Thermo Fisher Scientific), 50 μg/ml of RNase (Thermo Fisher Scientific) and 50 μg/mL lipase from *Aspergillus niger* (Sigma-Aldrich) in a buffer solution (20 mM Tris-HCl pH 7.0, 1 mM MgCl_2_, 0.05% sodium azide) for 4 h at 37 °C. The suspensions were washed with distilled water and treated with 200 μg/mL of trypsin (Sigma-Aldrich) in 20 mM Tris-HCl pH 8.0 overnight at 37°C with agitation. Finally, after inactivating trypsin (3-min boil), the suspensions were incubated with 48% hydrofluoric acid (Merck) overnight at 4 °C. After centrifugation (20 min, 25,000 × *g*), the pellet was washed twice with 250 mM Tris-HCl pH 7.0 and four times with distilled water to raise the pH to 5. The extracted peptidoglycan was lyophilized and resuspended in distilled water.

Muropeptides were prepared from purified peptidoglycan by overnight digestion with 2500 U/mL mutanolysin (Sigma-Aldrich) in 25 mM NaHPO_4_ pH 5.5, at 37°C with shaking. After reduction with sodium borohydride, muropeptide originating from peptidoglycan extracted from the same number of cells (1.5×10^9^) were analyzed by reverse phase-ultra high-pressure liquid chromatography (RP-UHPLC) using a 1290 chromatography system (Agilent Technologies) equipped with a Zorbax Eclipse Plus C18 RRHD column (100×2.1 mm, 1.8-μm particle size; Agilent Technologies). Elution was performed at 50 °C with 10 mM ammonium phosphate pH 5.6 and a linear gradient (0–20%, 270 min) of methanol, at a flow rate of 0.5 mL/min. Eluted muropeptides were detected by absorbance (202 nm).

### Gentamicin protection assay

JEG-3 and HepG2 cell lines were seeded in 24-well plates, with or without coverslips, to reach 90–100% confluency on the day of infection. Prior to seeding HepG2 cells, wells and coverslips were surface-coated with collagen (type I, rat tail) (Sigma-Aldrich) in a 50 µg/mL solution in Dulbecco’s phosphate-buffered saline (DPBS) (Gibco, Thermo Fisher Scientific) for 30 min and washed once with DPBS. On infection day, bacterial inocula were prepared by washing bacteria from overnight-grown, stationary-phase BHI cultures with DPBS and diluting them in serum-free medium. Cell monolayers were washed once with serum-free medium and infected for 1 h with the inocula at a multiplicity of infection (MOI) of 0.01 bacteria/JEG-3 cell or 5 bacteria/HepG2 cell. The inocula were removed from the wells and replaced with serum-supplemented medium containing 25 µg/mL of gentamicin (Sigma-Aldrich), to kill non-internalized bacteria. At 2 h, 6 h, 24 h and 72 h post-infection, cells were processed for immunofluorescence (see below) or for quantification of intracellular viable bacteria. In the latter case, cells were lysed in cold distilled water and serial dilutions of the lysates in DPBS were plated on BHI agar and incubated at 37 °C for at least 24 h for CFU enumeration.

### Generation of a polyclonal antiserum against CWD *Lm*

A rabbit polyclonal antiserum was raised against CWD *Lm* as follows. *Lm* 10403S were suspended in mineral water (10^8^ bacteria/mL), as described above, and incubated for 42 days. Bacteria were harvested by centrifugation (3,000 × *g*, 5 min), resuspended and incubated in fixative solution (1% (v/v) paraformaldehyde (PFA) in PBS) at 32 °C for 2 h, washed three times and resuspended in PBS. Bacterial neutralization was confirmed after plating on BHI agar and incubation at 37°C for several days.

Animal immunizations and serum recovery were performed by Covalab (Bron, France). White New Zealand female rabbits were inoculated with 1 mL of a 1:1 mixture of 10^8^ PFA-fixed CWD *Lm* and incomplete Freund’s adjuvant, and received boosts every three weeks for a total of three boosts. Immune serum was harvested at 53 and 74 days post-immunization and its reactivity and specificity towards CWD *Lm* was assessed by immunofluorescence microscopy.

### Immunofluorescence microscopy

Bacterial cells spotted onto poly-L-lysine-treated coverslips or coverslip-attached eukaryotic cells were fixed in a 4% (v/v) paraformaldehyde (PFA) solution in PBS for 20 or 30 min, respectively. Cells were washed in PBS, incubated in a blocking solution (2% bovine serum albumin in PBS) for 20 min and, in the case of eukaryotic cells, permeabilized (in a 0.4% (v/v) Triton X-100 solution in PBS for 4 min, followed by three washes in PBS) before proceeding with fluorescent labeling.

Rabbit *Listeria* O Antiserum Poly (anti-*Lm*; #223021, BD Difco) was used to label the *Listeria* CW. Oregon Green 488- or TRITC-conjugated WGA (Molecular Probes, Thermo Fisher Scientific) was used (25 µg/mL) to label the CW of serogroup 1/2 *Lm* strains (EGDe, 10403S). Rabbit anti-CWD *Lm* antiserum was used to label the exposed protoplast membrane of CW-shedding or CWD *Listeria*. Secondary antibodies consisted of goat and alpaca anti-rabbit antibodies conjugated with Alexa Fluor 488 (Molecular Probes, Thermo Fisher Scientific), Cy3 or Cy5 (Jackson ImmunoResearch). Alexa Fluor 647-conjugated phalloidin and Hoechst 33342 (Molecular Probes, Thermo Fisher Scientific) were respectively used to label F-actin and DNA, and, as with WGA, were added together with secondary antibodies. All incubations were made in blocking solution for 1 h in the dark.

Samples were mounted onto microscope glass slides with Fluoromount-G medium (Interchim) and examined on a ZEISS Axio Observer.Z1 epifluorescence microscope equipped with Plan-Apochromat 20×/0.8 NA (non-immersion), 40×/1.3 NA Oil and 100×/1.4 NA Oil (immersion) objectives, an Axiocam 506 Mono camera and operated with ZEN software (Carl Zeiss AG). Three to seven fields were acquired per coverslip and images were processed for quantification (see below) and/or figure montage with Fiji software.

### Image quantifications

Bacterial cell morphology was analyzed using Fiji software as follows: automatic thresholding was applied to phase-contrast images to select objects, and particle length and roundness parameters were selected (“Fit ellipse” and “Shape descriptors” options in “Set Measurements” menu) to be measured on thresholded objects (“Limit to threshold” option in “Set Measurements” menu). Outlier objects (i.e. too small/big, irregularly shaped) were excluded from the particle analysis (size: 0.5–1.5 µm^2^, circularity: 0.1–1.0). The length and roundness values per measured object were retrieved from the results table under the “Major” and “Round” columns, respectively.

To quantify the fraction of bacterial populations with CW and/or exposed plasma membrane (i.e. single or double) labeling, phase-contrast channel images were thresholded to select phase-contrast dense objects. Object outlines were then first laid over the DNA fluorescence channel to select bacterial cells, and afterwards over the fluorescence channels associated with CW and/or exposed plasma membrane to enumerate bacteria with single/double labeling. As co-labeling with the anti-*Lm* and anti-CWD *Lm* antibodies was not possible (same host species), only single-labeling quantifications were performed from separately labeled samples of the same bacterial population.

### Cryo-electron tomography

A solution of bovine serum albumin-coated gold tracer containing 10-nm colloidal gold fiducial particles (Aurion) was mixed with bacterial suspensions at a 2:1 ratio. This mixture was applied to the front (3.7 µL) and to the back (1.3 µL) of carbon-coated copper grids (R2/2, Cu 200 mesh; Quantifoil) previously glow-discharged (2 mA, 1.8×10^−1^ mbar, 1 min) in an ELMO system (Cordouan Technologies). Excess liquid was removed by blotting the grid backside with filter paper (9 sec, 18 °C, 95% humidity) and the sample was immediately frozen in liquid ethane in an EM GP automatic plunge freezer (Leica Microsystems). Grids were stored in liquid nitrogen until image acquisition.

Tilt series were acquired in a 300 kV Titan Krios G3 transmission electron microscope, equipped with a Cold FEG tip, a Selectris X energy filter with slit width set to 20 eV, single-tilt axis holder and a Falcon 4i direct electron detector (Thermo Fisher Scientific), and operated with the SerialEM software (version 4.0.13, U. Colorado Boulder, USA) (Mastronarde, 2003). Tilt series acquisition was performed in batches using a dose-symmetric scheme. One batch was obtained with an angular range of ±42° (3° increment), a defocus range of −3 to −8 µm, a pixel size of 4.8 Å (26,000x magnification), an exposure time of 8 s, a dose rate of 13.7 e/pixel/s and a total electron dose of about 140 e/Å^2^. Another batch was acquired with an angular range of ±50° (2° increment), a defocus range of −3 to −8 µm, a pixel size of 6.4 Å (19,500x magnification), an exposure time of 10 s, a dose rate of 12 e/pixel/s and a total electron dose of about 150 e/Å^2^. Tilt series were saved as separate stacks of frames, motion-corrected and restacked in order using the *alignframes* module in SerialEM.

3D reconstructions of tomograms were calculated in IMOD software (version 4.9.10, U. Colorado Boulder, USA) (Mastronarde & Held, 2017) by weighted back projections with dose weighting and a SIRT-like filter. The IMOD drawing tools and interpolator module were used to manually trace and produce a 3D surface of the bacterial plasma membrane. This surface was then imported to ChimeraX software (version 1.6, UC San Francisco, USA) (Pettersen et al., 2021) and used as a mask to extract slabs of subvolumes corresponding to the bacterial plasma membrane and CW. For the plasma membrane subvolume, the slab was produced using the *volume onesmask* function. For the CW, the *volume mask* function was used instead, cropping a larger slab beyond the CW limits, and the subvolume was visualized using the isosurface representation with an appropriate intensity threshold. Final visualizations and rendering were performed in ChimeraX using homemade scripts for video production.

### Fatty acid extraction and GC-MS analysis

Bacteria were harvested by centrifugation (3000 × *g*, 5 min), flash-frozen in liquid nitrogen and stored at −80 °C until further processing. Extraction and methylation of fatty acids (FA) were carried out directly on bacterial pellets as described (Touche et al., 2023). Whole-cell FA were first saponified and esterified by methanolic NaOH (1 mL of 3.75 M NaOH in 50% (v/v) methanol for 30 min at 100 °C) followed by methanolic HCl (addition of 2 mL of 3.25 M HCl in 45% (v/v) methanol solution and incubation for 10 min at 80°C). FA methyl esters (FAME) were then extracted with a 1:1 (v/v) diethyl ether/cyclohexane solution, and the organic phase was washed with dilute base (0.3 M NaOH).

Analytical gas chromatography of FAME was carried out in a GC-MS Trace 1300 / ISQ 7000 system (Thermo Fisher Scientific) equipped with a BPX70 capillary column (25 m, 0.22-mm internal diameter) (SGE, Victoria, Australia). Column temperature was set at 100 °C for 1 min and then increased to 170 °C at a rate of 2 °C/min. FA species were identified using MS databases (Replib, Mainlib, FAME2011). The relative abundance of FA species was expressed as the percentage of the total FAME peak area. Identified FA species were grouped in the following classes: iso and anteiso branched-chain FA (i-BFA and ai-BFA), saturated FA (SFA), and unsaturated FA (UFA).

### Laurdan generalized polarization

The generalized polarization of the lipophilic dye laurdan (6-dodecanoyl-2-dimethylaminonaphthalene), when bound to the *Lm* plasma membrane, was used as a measure of the bacterial membrane fluidity (Scheinpflug et al., 2017). Bacteria were sampled (1 mL) from mineral water suspensions (10^8^ cells/mL) at 0, 7, 14 and 28 days and incubated for 10 min in the dark with 10 µM of laurdan (Sigma-Aldrich) from a 1 mM stock in dimethylformamide (DMF). Unbound laurdan was washed off of bacterial cells with four cycles of centrifugation (8,000 × *g*, 5 min) and resuspension (vortex) in 1% (v/v) DMF in mineral water. After a final resuspension (vortex) in 1 mL of the washing solution, technical replicates (200 µL) were added to a clear-bottom black 96-well plate, which was then equilibrated to 25 °C in a Spark microplate reader (Tecan). After equilibration, laurdan fluorescence emission was induced at 350 nm and recorded at 450 and 500 nm. The generalized polarization (GP) of laurdan was determined by the formula: GP = (*I*_450_−*I*_500_)/(*I*_450_+*I*_500_), where *I* corresponds to the fluorescence intensity value at the recorded emission wavelength (Scheinpflug et al., 2017). The GP values obtained for every timepoint were normalized to those obtained from bacterial suspensions prepared on the same day (i.e. day 0). An increase of normalized laurdan GP values can be interpreted as a reduction of plasma membrane fluidity.

### Total internal reflection fluorescence correlation spectroscopy (TIR-FCS)

The diffusion of the lipophilic dye Nile red in the membrane of *Lm* was measured using total internal reflection fluorescence correlation spectroscopy (TIR-FCS) (Barbotin et al., 2023). In short, bacterial cells were labeled for 10 min with 0.1 µg/mL of Nile red from a 50 µg/mL stock in DMSO. FCS acquisitions were performed with a ZEISS Elyra PS1 TIRF microscope equipped with a Plan-Apochromat 100×/1.46 NA Oil immersion objective (Carl Zeiss AG). Fluorescence excitation at 561 nm was set to a power of ∼70 nW/µm². Each FCS acquisition consisted of a stack of 50,000 frames with a frame acquisition time of 1.26 ms, maximized by the use of only 10 lines of the camera chip. Pixels were binned 2 by 2 to increase signal levels to an effective pixel size of 320 nm. The resulting intensity timetraces were correlated and fitted as described (Barbotin et al., 2023).

TIR-FCS estimation of diffusion coefficients in small cells, such as bacteria, is biased by the cell morphology (Barbotin et al., 2023). To correct for this bias, the cell width and length of Nile red-stained bacteria were measured in epifluorescence images, using ImageJ. The average cell morphology values in each condition were then used to simulate TIR-FCS experiments to determine the diffusion coefficient bias (Barbotin et al., 2023). The corrected diffusion coefficient was obtained by dividing the experimentally measured diffusion coefficient value by the corresponding bias value.

### RNA extraction, RNA sequencing and gene set enrichment analysis

Bacteria (10^9^ cells) were harvested (5000 × *g*, 3 min) from mineral water suspensions (biological triplicates at 10^8^ cells/mL) in the first day and after 7 days of incubation, and immediately flash-frozen in liquid nitrogen and stored at −80 °C until further processing. Total RNAs were recovered from bead-beaten bacterial cells using a phenol/chloroform extraction method. Purified RNA samples were further prepared for sequencing at the I2BC sequencing platform (Gif-sur-Yvette, France)

RNA sample quality was assessed in an Agilent Bioanalyzer 2100, using the RNA 6000 pico kit (Agilent Technologies). Total RNAs (450 ng) were treated with DNase (Baseline-ZERO DNase, Epicentre) and ribosomal RNA was removed using the Illumina Ribo-Zero Magnetic Kit (Bacteria), according to the manufacturer recommendations. Directional RNA sequencing libraries were constructed using the TruSeq Stranded Total RNA Library Prep kit (Illumina) and sequenced (paired-end 2×75-bp) in a NextSeq500 instrument (Illumina). Alignment to the reference genome sequence of *Listeria monocytogenes* EGDe (RefSeq: NC_003210.1) was done using Bowtie2 (v2.4.4) (Langmead & Salzberg, 2012). For the detection of differentially expressed genes (DEGs), relative library sizes, fold changes (log_2_FC) and p-values were estimated using the R package “DESeq2” (v1.38.3) (Love et al., 2014), and p-values were then converted to q-values using the R package “fdrtool” (v1.2.17) (Strimmer, 2008). Genes with q-value ≤ 0.05 and absolute log_2_FC ≥ 1 were considered as DEGs.

Gene set enrichment analysis was performed on DEGs using the FUNAGE-Pro web server (http://funagepro.molgenrug.nl) (de Jong et al., 2022). A single-list analysis (gene locus tags) was done against the *Listeria monocytogenes* EGD-e reference genome (RefSeq: NC_003210.1). Gene ontology (GO) and KEGG pathway terms with a p-value ≤ 0.05 were considered to be statistically enriched.

### Chicken embryo infection assay

The chicken embryo model was used to test the recovery of *Lm* from a VBNC state in mineral water suspensions, as previously described (Andersson et al., 2015; Jean Michel Cappelier et al., 2007). Embryonated eggs from white Leghorn chickens raised under specific pathogen-free (SPF) conditions were obtained from the Infectiology of Farm, Model and Wildlife Animals Facility (PFIE) at the INRAE Centre Val de Loire (Nouzilly, France). Intact eggs were placed in an incubator (FIEM; Guanzate, Italy) at 37.7 °C and 47% humidity, under gentle rocking motion, to initiate embryo development. After 6 days, eggs were candled to check for signs of developing embryos, such as a strong vascularized network and embryo movement. Eggs showing underdeveloped or collapsed blood vessels were discarded, while eggs with no signs of embryo development (absence of blood vessels) were set aside to be used as “non-embryonated” eggs. Egg shells were sterilized with 70% ethanol and perforated just above the border of the air sac to allow the injection of 100 µL of bacterial suspension into the allantoic cavity (or albumen in non-embryonated eggs), using a 25G (0.5×16 mm) needle. Shell punctures were sealed with a sticky tag and the eggs were returned to the incubator. At 48 h post-inoculation, embryonated eggs were candled to discard dead embryos, and viable embryos were euthanized by incubation at 4 °C for 2 h. Embryos were recovered in aseptic conditions and placed into a tube with 4 mL of sterile DPBS to undergo mechanical homogenization (T-25 Ultra-Turrax, IKA). Serial dilutions of the embryo homogenate (or albumen from non-embryonated eggs) in DPBS were plated (500 µL) on BHI agar and incubated at 37 °C for at least 24 h to assess the presence of culturable *Lm*.

For inoculation of eggs with VBNC *Lm*, suspensions of EGDe-GFP at 10^6^ bacteria/mL were prepared 28 days before, and their viability and culturability were checked to select the one with the lowest residual culturability. As a result, we used an inoculum containing 10^6^ viable *Lm*/mL and 0.5 *Lm* CFU/mL, which corresponded to an inoculated dose containing 10^5^ viable and 0.05 culturable bacteria. Injections of 100 µL of mineral water or a suspension of vegetative EGDe-GFP – prepared by washing and diluting bacteria from an overnight-grown culture to approximately 5,000 CFU/mL – were included as negative and positive controls, respectively.

To assess whether bacterial growth recovered from inoculated eggs resulted from the revival of VBNC cells or solely from the regrowth of residual culturable bacteria, we compared the frequency of bacterial growth obtained before and after egg inoculation with the VBNC *Lm* suspension. The frequency before inoculation was determined by serially inoculating wells of a 96-well plate containing 100 µL of BHI broth with 100 µL of the VBNC *Lm* suspension, and calculating the fraction of inoculated wells showing bacterial growth after 48 h of incubation at 37 °C. Similarly, the frequency after inoculation was determined from the fraction of inoculated eggs scored positive for bacterial growth on BHI agar.

### Statistics

Data from experiments comprising biological replicates (n=3) are graphically presented as mean ± standard deviation or median ± interquartile range. Prism software (version 9.0; GraphPad) was used to generate graphs and to perform statistical analyses (tests reported in figure legends). Differences between means were considered statistically significant for p values ≤ 0.05.

## Data availability

The RNA sequencing data generated in this study have been deposited in NCBI’s Gene Expression Omnibus and are accessible through GEO Series accession number GSE246157 (https://www.ncbi.nlm.nih.gov/geo/query/acc.cgi?acc=GSE246157). Additional data and material generated by this work are available upon request.

## Acknowledgements

We thank the members of the EpiMic team for helpful discussions. We are grateful to Didier Cabanes (U. Porto, Portugal), for kindly gifting us the EGDeΔ*ami* and EGDeΔ*aut* strains; to Daniel Portnoy (UC Berkeley, California, USA), for the 10403SΔ*p60*, 10403SΔ*rpf1-2* and 10403SΔ*relAPQ* strains; and to Anna Oevermann (U. Bern, Switzerland), for the JF5203 strain. We also thank Jasmina Vidic, for help with the laurdan GP assays; Matthieu Bertrand, for assistance with fluorescence microscopy image quantification scripts; and to Simonetta Gribaldo, Delphine Lechardeur, Philippe Noirot and Pascale Cossart, for their critical feedback to this manuscript. We acknowledge the sequencing and bioinformatics expertise of the I2BC high-throughput sequencing facility, supported by France Génomique (funded by the French government’s Programme Investissements d’Avenir (PIA): ANR-10-INBS-0009). We also acknowledge the cryo-ET expertise and assistance of the Institut Pasteur’s NanoImaging Core facility, created and supported by a PIA grant (EquipEx CACSICE: ANR-11-EQPX-0008).

This work was supported by grants from the Agence Nationale de la Recherche (ANR) to HB (PERMALI: ANR-20-CE35-0001-01) and AP (THOR: ANR-20-CE15-0008-01); from the Université Paris-Saclay to HB (DEPISTALIS, AAP Poc in labs 2019); from the INRAE’s MICA department and Micalis Institute to AP (AAP Micalis 2023); and from the European Research Council, under the Horizon 2020 research and innovation program, to RCL (agreement ID: 772178). FC was supported by postdoctoral grants from INRAE MICA department and ANR (ANR-20-PAMR-0011). AB received funding from the European Union’s Horizon 2020 research and innovation program, under the Marie Skłodowska-Curie Actions grant (agreement ID: 101030628).

We dedicate this work to the memory of Hélène Bierne and Fabrizia Stavru.

## Author contributions

FC, HB and AP conceived the work plan and methodology. EM conceived and performed pilot experiments. FC, AC, ASR, ST, ADG, PC, PN, FDB, AB, ED, KG, CS, MPCC, EM and AP performed experiments. FC, ASR, ADG, PN, FDB and AB analyzed collected data. ASR, ST, PN, FDB, KG, RCL, MPCC, HB and AP provided resources (reagents, equipment and/or analysis tools). FC, HB and AP wrote the original draft. FC, HB and AP reviewed and edited the draft. FC prepared and organized data visualization. HB and AP provided funding and supervision. All authors contributed to this work and approved the submitted version.

## Conflicts of interest

The authors declare no conflict of interest.

## Extended Data figure legends

**Extended Data Fig. 1.**
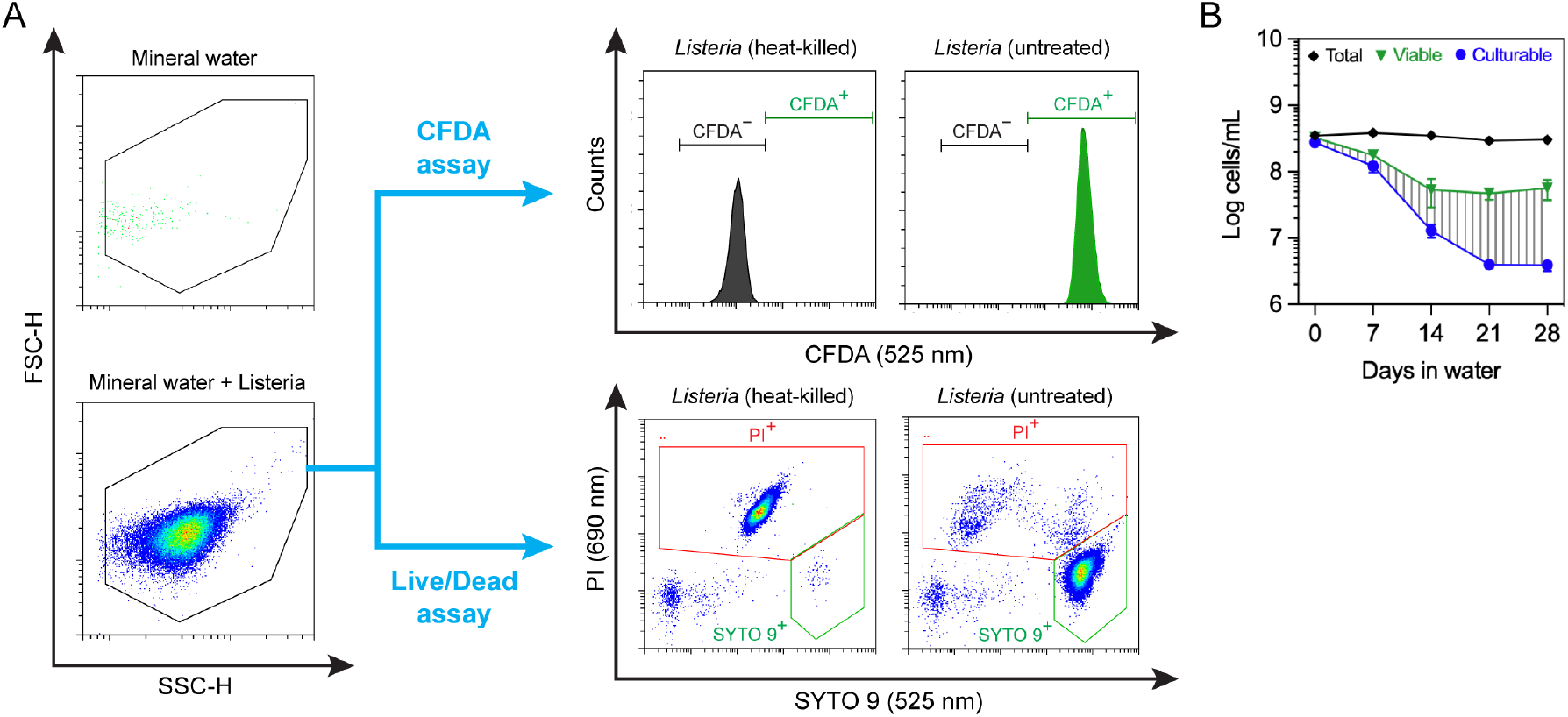
Quantification of the viable cell populations by flow cytometry. **(A)** Schematic diagram of the flow cytometry protocol (detection parameters and gating) applied for enumeration of viable bacterial cells in mineral water suspensions using the CFDA and Live/Dead viability assays. **(B)** Representation of the data depicted in Fig. 1A, showing an alternative quantification of viable cell population using the Live/Dead assay. The area covered with vertical grey dashes indicates the VBNC cell population.

**Extended Data Fig. 2.**
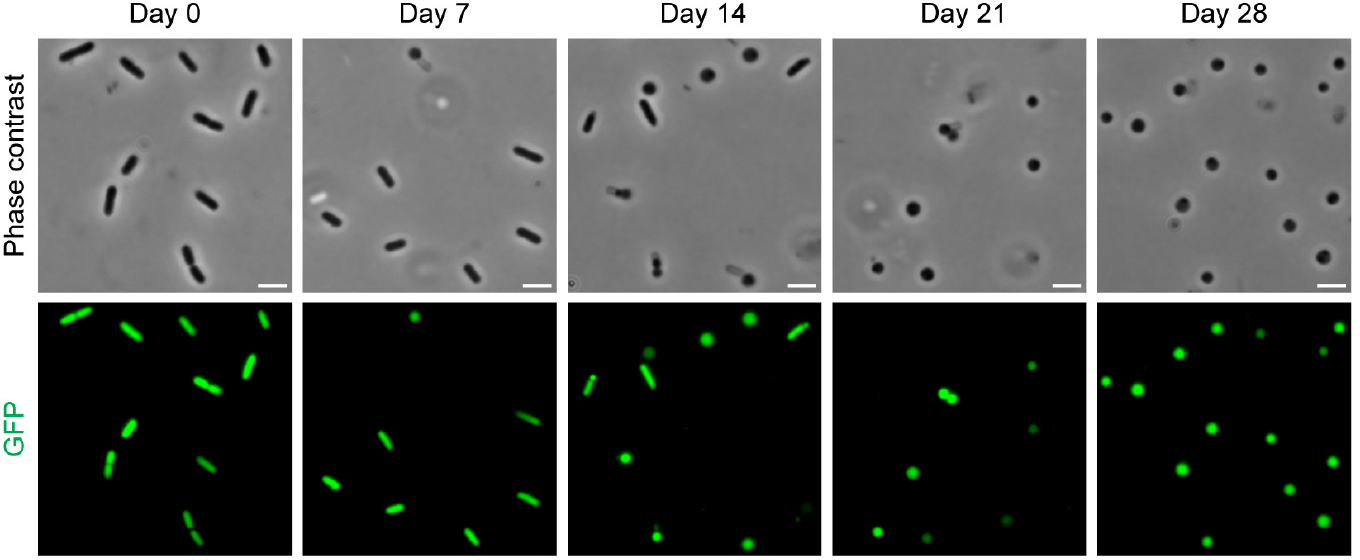
Coccoid cell forms derive from rod-shaped *Lm*. Phase-contrast and fluorescence micrographs of GFP-expressing *Lm* EGDe sampled from mineral water suspensions (10^8^ cells/mL) at the indicated timepoints. Scale bar: 2 µm.

**Extended Data Fig. 3.**
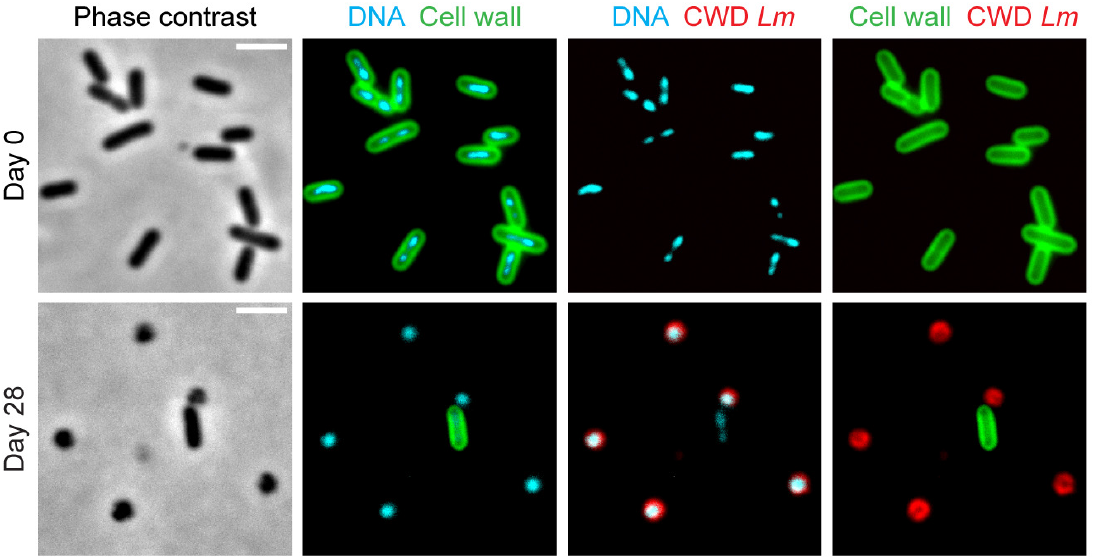
A polyclonal antiserum raised against CWD VBNC *Lm* labels specifically bacteria with externally exposed membrane. Phase-contrast and fluorescence micrographs of *Lm* EGDe sampled from mineral water suspensions at the indicated timepoints. Bacteria were fixed and fluorescently labelled for visualization of DNA (Hoechst 33342) and CW (WGA), and to assess the specificity of the rabbit polyclonal antiserum raised against CWD *Lm* (anti-CWD *Lm*). Scale bar: 2 µm.

**Extended Data Fig. 4.**
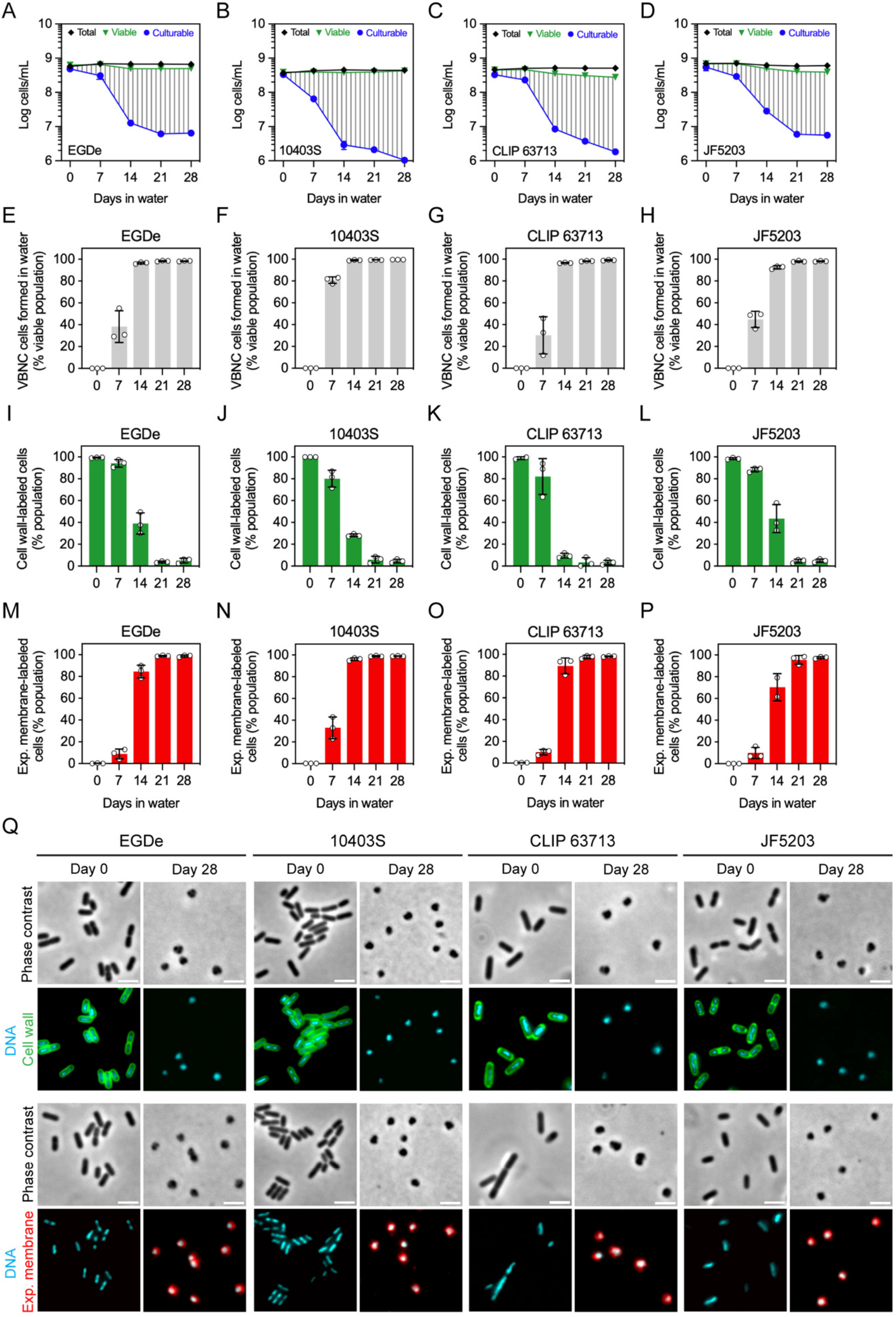
Various *Lm* strains are able to reach a CWD VBNC state. **(A–D)** Total, viable and culturable cell profiles of mineral water suspensions of *Lm* strains EGDe (A), 10403S (B), CLIP 63713 (C) and JF5203 (D). Culturable bacteria were determined by enumeration of CFUs, while total and viable bacteria were quantified by flow cytometry. Data are represented as the mean ± SD from three independent suspensions. The area covered with vertical grey dashes indicates the VBNC cell population. **(E–H)** Fraction of the EGDe (E), 10403S (F), CLIP 63713 (G) and JF5203 (H) populations consisting of VBNC cells formed in mineral water. Data are represented as the mean ± SD from three independent suspensions, each indicated by a dot. **(I–P)** Fraction of the EGDe (I, M), 10403S (J, N), CLIP 63713 (K, O) and JF5203 (L, P) populations showing CW (I–L) or exposed membrane (M–P) labelling by fluorescence microscopy, at the indicated timepoints. Data are represented as the mean ± SD from three independent suspensions, each indicated by a dot. **(Q)** Phase-contrast and fluorescence micrographs of EGDe, 10403S, CLIP 63713 and JF5203 sampled from mineral water suspensions at the indicated timepoints. Bacteria were fixed and fluorescently labelled for visualization of DNA (Hoechst 33342), CW (anti-*Lm*) and exposed plasma membrane (anti-CWD *Lm*). Scale bar: 2 µm.

**Extended Data Fig. 5.**
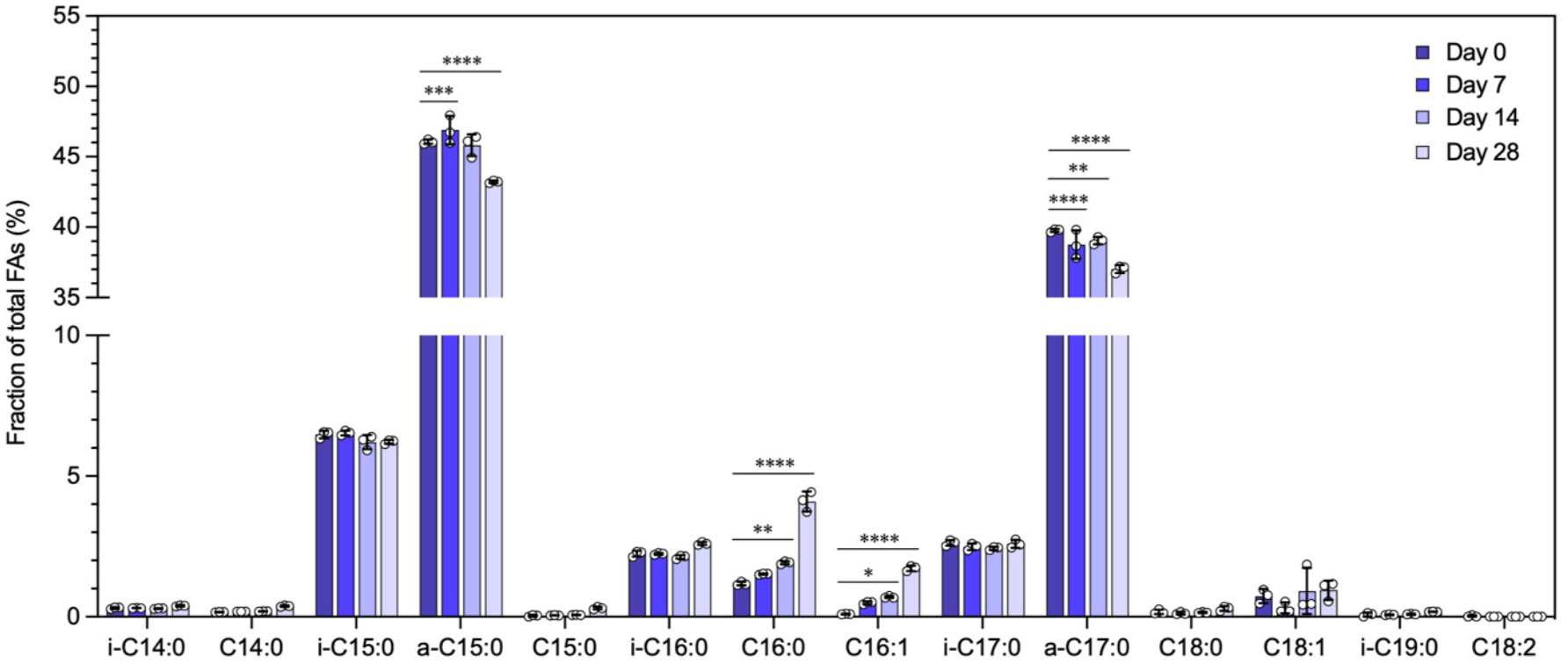
Changes in the fatty acid composition of the *Lm* plasma membrane during incubation in mineral water. The fatty acid (FA) composition of the plasma membrane of *Lm* EGDe sampled from mineral water suspensions at the indicated timepoints was determined by gas chromatography-mass spectrometry (GC–MS/MS). Chain length and saturation of identified FA species is indicated in the x-axis by C#:$, where # corresponds to the number of carbons and $ the number of double bonds in the chain. Branched-chain FA species are indicated by the prefixes i-(iso) and a-(anteiso). Data are represented as the mean (bar) ± SD from three independent suspensions, each indicated by a dot. Statistical analysis was performed using a two-way ANOVA test with Dunnett’s post hoc multiple comparison correction. *, p≤0.05; **, p≤0.01; ***, p≤0.001; ****, p≤0.0001.

**Extended Data Fig. 6.**
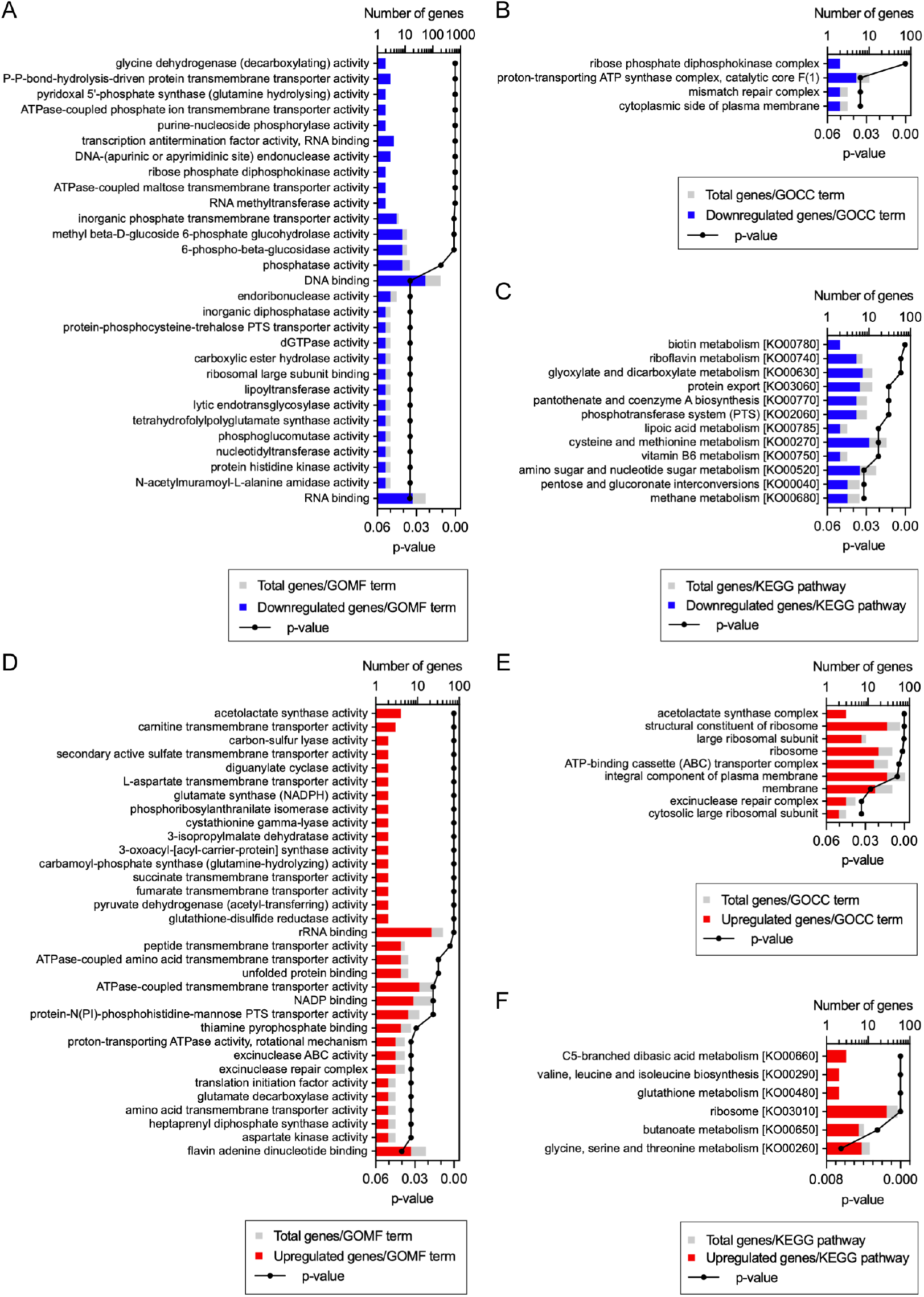
Extended functional enrichment analysis of differentially regulated genes. **(A–F)** Downregulated (A–C) and upregulated (D–F) *Lm* EGDe genes analyzed for statistically overrepresented gene ontology molecular function (A, D) and cellular component (GOCC) terms (B, E), and KEGG pathways (C, F). Terms and pathways are ranked from top to bottom by increasing p-value and decreasing fraction of differentially regulated genes per category/pathway.

**Extended Data Fig. 7.**
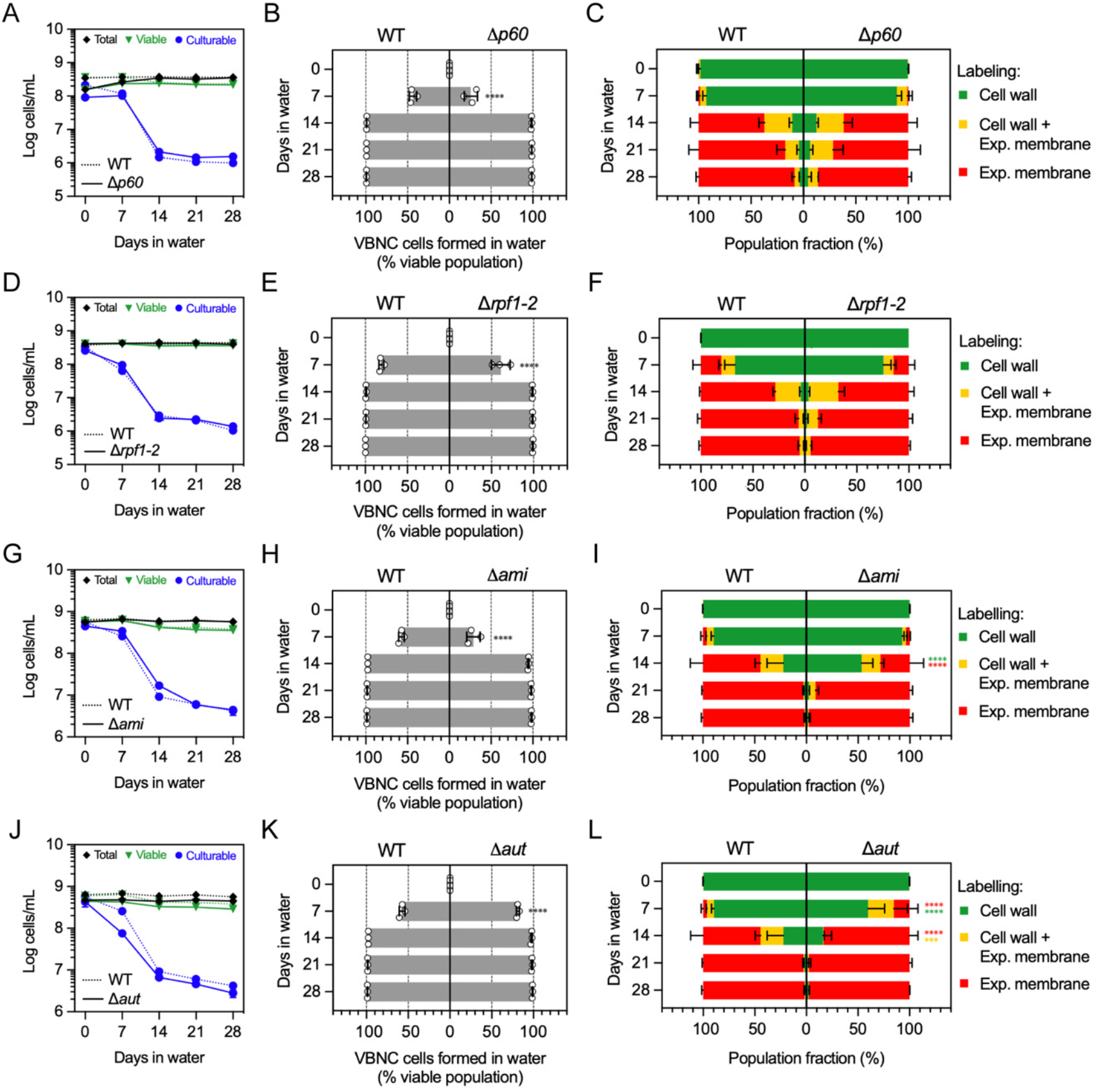
The *Lm* autolysins p60, Rpf1/2, Ami and Auto are not required for CW loss and VBNC state transition. **(A, D, G, J)** Total, viable and culturable cell profiles of mineral water suspensions of wild type (WT) and isogenic *Lm* mutant strains lacking p60 (A), Rpf1 and Rpf2 (D), Ami (G), or Auto (J) autolysins. Culturable bacteria were determined by enumeration of CFUs, while total and viable bacteria were quantified by flow cytometry. Data are represented as the mean ± SD from three independent suspensions. **(B, E, H, K)** Fraction of the WT and Δ*p60* (B), Δ*rpf1-2* (E), Δ*ami* (H) or Δ*aut* (K) populations consisting of VBNC cells formed in mineral water. Data are represented as the mean (bar) ± SD from three independent suspensions, each indicated by a dot. Statistical analysis between mutant and WT strains was performed using a two-way ANOVA test with Šidák’s post hoc multiple comparison correction. ****, p≤0.0001. **(C, F, I, L)** Fraction of the WT and Δ*p60* (C), Δ*rpf1-2* (F), Δ*ami* (I) or Δ*aut* (L) populations showing either one or both CW and exposed plasma membrane labelling by fluorescence microscopy, at the indicated timepoints. Data are represented as stacked bars (one for each labelling group) with mean ± SD from three independent suspensions. Statistical analysis between mutant and WT strains was performed for each labelling group using a two-way ANOVA test with Tukey’s post hoc multiple comparison correction. *, p≤0.05; ***, p≤0.001; ****, p≤0.0001.

**Supplementary Table 1.** Composition of mineral water used in this study.

| Component | mg/L |
| --- | --- |
| Bicarbonates ($\text{HCO}_3^-$) | 74 |
| Calcium ($\text{Ca}^{2+}$) | 12 |
| Chlorides ($\text{Cl}^-$) | 15 |
| Magnesium ($\text{Mg}^{2+}$) | 8 |
| Nitrates ($\text{NO}_3^-$) | 7.3 |
| Potassium ($\text{K}^+$) | 6 |
| Silica ($\text{SiO}_2$) | 32 |
| Sodium ($\text{Na}^+$) | 12 |
| Sulfates ($\text{SO}_4^{2-}$) | 9 |
| Total dry residue (180 °C) | 130 |

**Supplementary Table 6.** VBNC *Lm* revert back to a culturable state after passage in embryonated chicken eggs.

| Inoculum | Culturability before egg passage <sup>(1)</sup> | Culturability after egg passage <sup>(2)</sup> |  |  |  |
| --- | --- | --- | --- | --- | --- |
|  |  | Embryonated eggs | Fisher's exact test (p-value) <sup>(3)</sup> | Non-embryonated eggs | Fisher's exact test (p-value) <sup>(4)</sup> |
| Mineral water | 0/3 (0%) | 0/10 (0%) | >0.999 | 0/10 (0%) | >0.999 |
| VBNC *Lm* | 8/84 (9.52%) | 24/24 (100%) | $1.63 \times 10^{-17}$ | 0/18 (0%) | 0.344 |
| Vegetative *Lm* | 3/3 (100%) | 8/8 (100%) | >0.999 | 2/2 (100%) | >0.999 |

<sup>(1)</sup> Number of BHI wells with bacterial growth / number of BHI wells inoculated
<sup>(2)</sup> Number of eggs with bacterial growth / number of eggs inoculated
<sup>(3)</sup> Comparison of culturability before egg passage and after passage in embryonated eggs
<sup>(4)</sup> Comparison of culturability before egg passage and after passage in non-embryonated eggs

**Supplementary Table 7.**
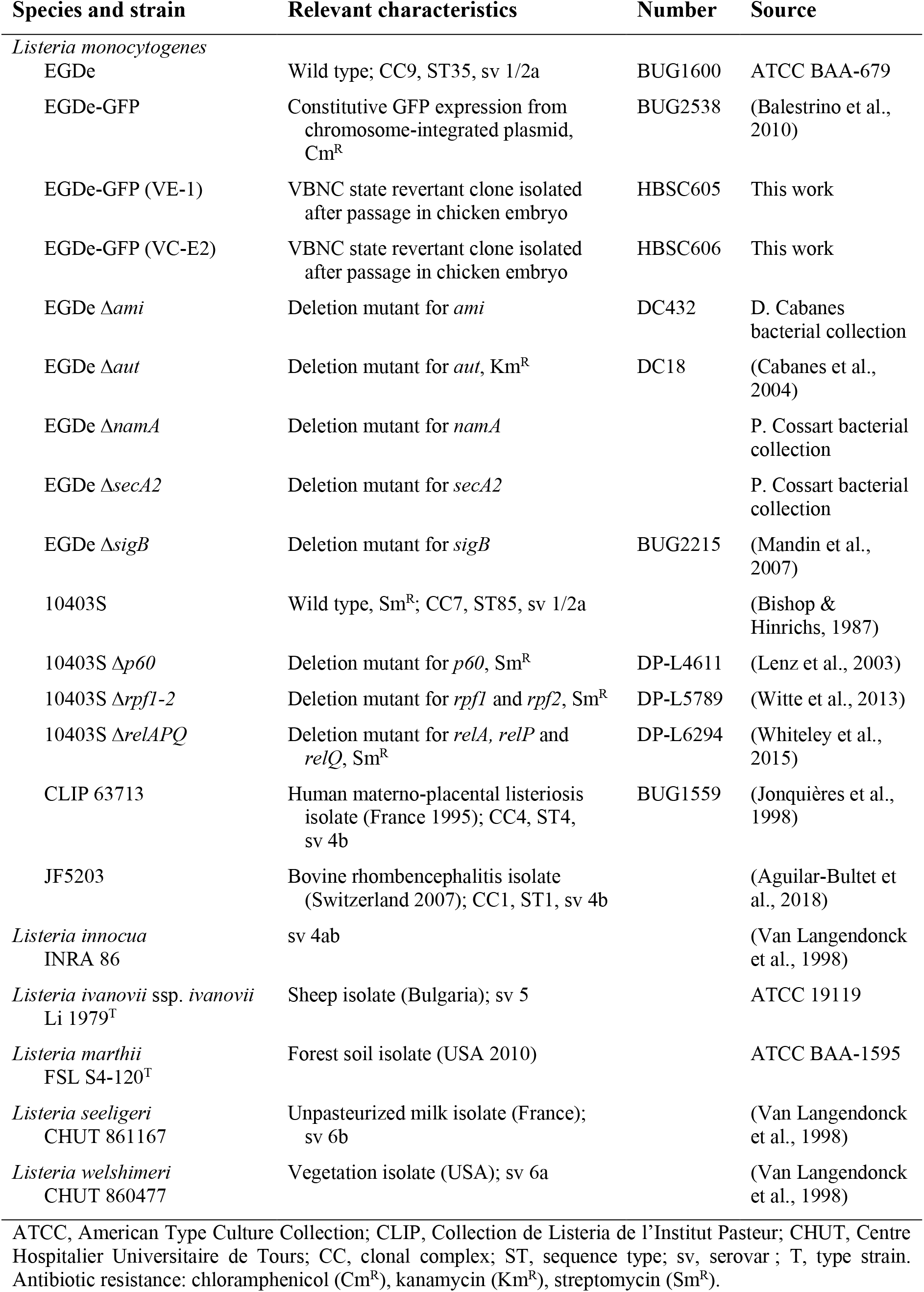
Bacteria used in this study.

## Notes

### Competing Interest Statement

The authors have declared no competing interest.

## References

Aguilar-Bultet, L., Nicholson, P., Rychener, L., Dreyer, M., Gözel, B., Origgi, F. C., Oevermann, A., Frey, J., & Falquet, L. (2018). Genetic Separation of Listeria monocytogenes Causing Central Nervous System Infections in Animals. Frontiers in Cellular and Infection Microbiology, 8(FEB), 20. 10.3389/fcimb.2018.00020

Andersson, C., Gripenland, J., & Johansson, J. (2015). Using the chicken embryo to assess virulence of Listeria monocytogenes and to model other microbial infections. Nature Protocols, 10(8), 1155–1164. 10.1038/nprot.2015.073

Argov, T., Sapir, S. R., Pasechnek, A., Azulay, G., Stadnyuk, O., Rabinovich, L., Sigal, N., Borovok, I., & Herskovits, A. A. (2019). Coordination of cohabiting phage elements supports bacteria–phage cooperation. Nature Communications 2019 10:1, 10(1), 1–14. 10.1038/s41467-019-13296-x

Ayrapetyan, M., Williams, T., & Oliver, J. D. (2018). Relationship between the Viable but Nonculturable State and Antibiotic Persister Cells. Journal of Bacteriology, 200(20), 580. 10.1128/JB.00249-18

Bai, K., Yan, H., Chen, X., Lyu, Q., Jiang, N., Li, J., & Luo, L. (2021). The Role of RelA and SpoT on ppGpp Production, Stress Response, Growth Regulation, and Pathogenicity in Xanthomonas campestris pv. campestris. Microbiology Spectrum, 9(3). 10.1128/SPECTRUM.02057-21/SUPPL_FILE/SPECTRUM02057-21_SUPP_1_SEQ9.PDF

Baker, R. M., Singleton, F. L., & Hood, M. A. (1983). Effects of nutrient deprivation on Vibrio cholerae. Applied and Environmental Microbiology, 46(4), 930–940. 10.1128/aem.46.4.930-940.1983

Balestrino, D., Hamon, M. A., Dortet, L., Nahori, M.-A. A., Pizarro-Cerda, J., Alignani, D., Dussurget, O., Cossart, P., Toledo-Arana, A., Anne Hamon, M., Dortet, L., Nahori, M.-A. A., Pizarro-Cerda, J., Alignani, D., Dussurget, O., Cossart, P., & Toledo-Arana, A. (2010). Single-cell techniques using chromosomally tagged fluorescent bacteria to study Listeria monocytogenes infection processes. Applied and Environmental Microbiology, 76(11), 3625–3636. 10.1128/AEM.02612-09

Barbotin, A., Billaudeau, C., Sezgin, E., & Lopez, R. C. (2023). Quantification of membrane fluidity in bacteria using TIR-FCS. BioRxiv, 2023.10.13.562271. 10.1101/2023.10.13.562271

Beskrovnaya, P., Sexton, D. L., Golmohammadzadeh, M., Hashimi, A., & Tocheva, E. I. (2021). Structural, Metabolic and Evolutionary Comparison of Bacterial Endospore and Exospore Formation. Frontiers in Microbiology, 12, 452. 10.3389/FMICB.2021.630573/BIBTEX

Besnard, V., Federighi, M., & Cappelier, J. M. (2000a). Development of a direct viable count procedure for the investigation of VBNC state in Listeria monocytogenes. Letters in Applied Microbiology, 31(1), 77–81. 10.1046/j.1472-765x.2000.00771.x

Besnard, V., Federighi, M., & Cappelier, J. M. (2000b). Evidence of Viable But Non-Culturable state in Listeria monocytogenes by direct viable count and CTC-DAPI double staining. Food Microbiology, 17(6), 697–704. 10.1006/fmic.2000.0366

Besnard, V., Federighi, M., Declerq, E., Jugiau, F., & Cappelier, J. M. (2002). Environmental and physico-chemical factors induce VBNC state in Listeria monocytogenes. Veterinary Research, 33(4), 359–370. 10.1051/vetres:2002022

Bierne, H., & Cossart, P. (2007). Listeria monocytogenes Surface Proteins: from Genome Predictions to Function. Microbiology and Molecular Biology Reviews, 71(2), 377–397. 10.1128/MMBR.00039-06

Bishop, D. K., & Hinrichs, D. J. (1987). Adoptive transfer of immunity to Listeria monocytogenes. The influence of in vitro stimulation on lymphocyte subset requirements. Journal of Immunology (Baltimore, Md. : 1950), 139(6), 2005–2009. 10.4049/JIMMUNOL.139.6.2005

Boaretti, M., Lleò, M. D. M., Bonato, B., Signoretto, C., & Canepari, P. (2003). Involvement of rpoS in the survival of Escherichia coli in the viable but non-culturable state. Environmental Microbiology, 5(10), 986–996. 10.1046/J.1462-2920.2003.00497.X

Bremer, P. J., Osborne, C. M., Kemp, R. A., & Smith, J. J. (1998). Survival of Listeria monocytogenes in sea water and effect of exposure on thermal resistance. Journal of Applied Microbiology, 85(3), 545–553. 10.1046/j.1365-2672.1998.853533.x

Cabanes, D., Dussurget, O., Dehoux, P., & Cossart, P. (2004). Auto, a surface associated autolysin of Listeria monocytogenes required for entry into eukaryotic cells and virulence. Molecular Microbiology, 51(6), 1601–1614. 10.1111/j.1365-2958.2003.03945.x

Cappelier, J. M., Minet, J., Magras, C., Colwell, R. R., & Federighi, M. (1999). Recovery in embryonated eggs of viable but nonculturable Campylobacter jejuni cells and maintenance of ability to adhere to HeLa cells after resuscitation. Applied and Environmental Microbiology, 65(11), 5154–5157. 10.1128/AEM.65.11.5154-5157.1999

Cappelier, Jean Michel, Besnard, V., Roche, S. M., Velge, P., & Federighi, M. (2007). Avirulent viable but non culturable cells of Listeria monocytogenes need the presence of an embryo to be recovered in egg yolk and regain virulence after recovery. Veterinary Research, 38(4), 573–583. 10.1051/vetres:2007017

Carroll, S. A., Hain, T., Technow, U., Darji, A., Pashalidis, P., Josep, S. W., & Chakraborty, T. (2003). Identification and Characterization of a Peptidoglycan Hydrolase, MurA, of Listeria monocytogenes, a Muramidase Needed for Cell Separation. Journal of Bacteriology, 185(23), 6801–6808. 10.1128/JB.185.23.6801-6808.2003

Chaveerach, P., ter Huurne, A. A. H. M., Lipman, L. J. A., & van Knapen, F. (2003). Survival and resuscitation of ten strains of Campylobacter jejuni and Campylobacter coli under acid conditions. Applied and Environmental Microbiology, 69(1), 711–714. 10.1128/AEM.69.1.711-714.2003

Claessen, D., & Errington, J. (2019). Cell Wall Deficiency as a Coping Strategy for Stress. Trends in Microbiology, 27(12), 1025–1033. 10.1016/j.tim.2019.07.008

Costa, K., Bacher, G., Allmaier, G., Dominguez-Bello, M. G., Engstrand, L., Falk, P., De Pedro, M. A., & García-del Portillo, F. (1999). The morphological transition of Helicobacter pylori cells from spiral to coccoid is preceded by a substantial modification of the cell wail. Journal of Bacteriology, 181(12), 3710–3715. 10.1128/JB.181.12.3710-3715.1999/ASSET/8E5217C5-674A-40E6-845D-DD583C0A0E41/ASSETS/GRAPHIC/JB1290033003.JPEG

Cunningham, E., O’Byrne, C., & Oliver, J. D. (2009). Effect of weak acids on Listeria monocytogenes survival: Evidence for a viable but nonculturable state in response to low pH. Food Control, 20(12), 1141–1144. 10.1016/j.foodcont.2009.03.005

Dannenberg, N., Bravo, V. C., Weijers, T., Spaink, H., Ottenhoff, T., Briegel, A., & Claessen, D. (2022). Mycobacteria form viable cell wall-deficient cells that are undetectable by conventional diagnostics. BioRxiv, 2022.11.16.516772. 10.1101/2022.11.16.516772

de Jong, A., Kuipers, O. P., & Kok, J. (2022). FUNAGE-Pro: comprehensive web server for gene set enrichment analysis of prokaryotes. Nucleic Acids Research, 50(W1), W330–W336. 10.1093/nar/gkac441

Dell’Era, S., Buchrieser, C., Couvé, E., Schnell, B., Briers, Y., Schuppler, M., & Loessner, M. J. (2009). Listeria monocytogenes l-forms respond to cell wall deficiency by modifying gene expression and the mode of division. Molecular Microbiology, 73(2), 306–322. 10.1111/j.1365-2958.2009.06774.x

Domínguez-Cuevas, P., Mercier, R., Leaver, M., Kawai, Y., & Errington, J. (2012). The rod to L-form transition of Bacillus subtilis is limited by a requirement for the protoplast to escape from the cell wall sacculus. Molecular Microbiology, 83(1), 52–66. 10.1111/j.1365-2958.2011.07920.x

Dong, K., Pan, H., Yang, D., Rao, L., Zhao, L., Wang, Y., & Liao, X. (2020). Induction, detection, formation, and resuscitation of viable but non-culturable state microorganisms. Comprehensive Reviews in Food Science and Food Safety, 19(1), 149–183. 10.1111/1541-4337.12513

Duru, I. C., Bucur, F. I., Andreevskaya, M., Nikparvar, B., Ylinen, A., Grigore-Gurgu, L., Rode, T. M., Crauwels, P., Laine, P., Paulin, L., Løvdal, T., Riedel, C. U., Bar, N., Borda, D., Nicolau, A. I., & Auvinen, P. (2021). High-pressure processing-induced transcriptome response during recovery of Listeria monocytogenes. BMC Genomics, 22(1), 1–20. 10.1186/S12864-021-07407-6/FIGURES/5

Errington, J., Mickiewicz, K., Kawai, Y., & Wu, L. J. (2016). L-form bacteria, chronic diseases and the origins of life. Philosophical Transactions of the Royal Society B: Biological Sciences, 371(1707), 20150494. 10.1098/rstb.2015.0494

Fiedler, F. (1988). Biochemistry of the cell surface of Listeria strains: A locating general view. Infection, 16(2 Supplement). 10.1007/BF01639729

Highmore, C. J., Warner, J. C., Rothwell, S. D., Wilks, S. A., & Keevil, C. W. (2018). Viable-but-Nonculturable Listeria monocytogenes and Salmonella enterica Serovar Thompson Induced by Chlorine Stress Remain Infectious. MBio, 9(2). 10.1128/mBio.00540-18

Höltje, J.-V. V. (1998). Growth of the Stress-Bearing and Shape-Maintaining Murein Sacculus of Escherichia coli. Microbiology and Molecular Biology Reviews : MMBR, 62(1), 181–203. 10.1128/mmbr.62.1.181-203.1998

Irving, S. E., Choudhury, N. R., & Corrigan, R. M. (2021). The stringent response and physiological roles of (pp)pGpp in bacteria. Nature Reviews. Microbiology, 19(4), 256–271. 10.1038/s41579-020-00470-y

Ivy, R. A., Wiedmann, M., & Boor, K. J. (2012). Listeria monocytogenes grown at 7°C shows reduced acid survival and an altered transcriptional response to acid shock compared to L. Monocytogenes grown at 37°C. Applied and Environmental Microbiology, 78(11), 3824–3836. 10.1128/AEM.00051-12/SUPPL_FILE/AEM-AEM00051-12-S08.PDF

Jonquières, R., Bierne, H., Mengaud, J., & Cossart, P. (1998). The inlA gene of Listeria monocytogenes LO28 harbors a nonsense mutation resulting in release of internalin. Infection and Immunity, 66(7), 3420–3422. 10.1128/IAI.66.7.3420-3422.1998

Kawai, Y., Mickiewicz, K., & Errington, J. (2018). Lysozyme Counteracts β-Lactam Antibiotics by Promoting the Emergence of L-Form Bacteria. Cell, 172(5), 1038–1049.e10. 10.1016/j.cell.2018.01.021

Langmead, B., & Salzberg, S. L. (2012). Fast gapped-read alignment with Bowtie 2. Nature Methods 2012 9:4, 9(4), 357–359. 10.1038/nmeth.1923

Lenz, L. L., Mohammadi, S., Geissler, A., & Portnoy, D. A. (2003). SecA2-dependent secretion of autolytic enzymes promotes Listeria monocytogenes pathogenesis. Proceedings of the National Academy of Sciences of the United States of America, 100(21), 12432–12437. 10.1073/pnas.2133653100

Li, L., Mendis, N., Trigui, H., Oliver, J. D., & Faucher, S. P. (2014). The importance of the viable but non-culturable state in human bacterial pathogens. Frontiers in Microbiology, 5(JUN), 258. 10.3389/fmicb.2014.00258

Lindbäck, T., Rottenberg, M. E., Roche, S. M., & Rørvik, L. M. (2010). The ability to enter into an avirulent viable but non-culturable (VBNC) form is widespread among Listeria monocytogenes isolates from salmon, patients and environment. Veterinary Research, 41(1), 8. 10.1051/vetres/2009056

Lotoux, A., Milohanic, E., & Bierne, H. (2022). The Viable But Non-Culturable State of Listeria monocytogenes in the One-Health Continuum. Frontiers in Cellular and Infection Microbiology, 12, 849915. 10.3389/fcimb.2022.849915

Love, M. I., Huber, W., & Anders, S. (2014). Moderated estimation of fold change and dispersion for RNA-seq data with DESeq2. Genome Biology, 15(12), 550. 10.1186/s13059-014-0550-8

Lyautey, E., Lapen, D. R., Wilkes, G., McCleary, K., Pagotto, F., Tyler, K., Hartmann, A., Piveteau, P., Rieu, A., Robertson, W. J., Medeiros, D. T., Edge, T. A., Gannon, V., & Topp, E. (2007). Distribution and characteristics of Listeria monocytogenes isolates from surface waters of the South Nation River watershed, Ontario, Canada. Applied and Environmental Microbiology, 73(17), 5401–5410. 10.1128/AEM.00354-07

Machata, S., Hain, T., Rohde, M., & Chakraborty, T. (2005). Simultaneous deficiency of both MurA and p60 proteins generates a rough phenotype in Listeria monocytogenes. Journal of Bacteriology, 187(24), 8385–8394. 10.1128/JB.187.24.8385-8394.2005

Mandin, P., Repoila, F., Vergassola, M., Geissmann, T., & Cossart, P. (2007). Identification of new noncoding RNAs in Listeria monocytogenes and prediction of mRNA targets. Nucleic Acids Research, 35(3), 962–974. 10.1093/nar/gkl1096

Mastronarde, D. N. (2003). SerialEM: A Program for Automated Tilt Series Acquisition on Tecnai Microscopes Using Prediction of Specimen Position. Microscopy and Microanalysis, 9(S02), 1182–1183. 10.1017/S1431927603445911

Mastronarde, D. N., & Held, S. R. (2017). Automated tilt series alignment and tomographic reconstruction in IMOD. Journal of Structural Biology, 197(2), 102–113. 10.1016/J.JSB.2016.07.011

Noll, M., Trunzer, K., Vondran, A., Vincze, S., Dieckmann, R., Al Dahouk, S., & Gold, C. (2020). Benzalkonium Chloride Induces a VBNC State in Listeria monocytogenes. Microorganisms, 8(2). 10.3390/microorganisms8020184

Parry, B. R., Surovtsev, I. V., Cabeen, M. T., O’Hern, C. S., Dufresne, E. R., & Jacobs-Wagner, C. (2014). The bacterial cytoplasm has glass-like properties and is fluidized by metabolic activity. Cell, 156(1–2), 183–194. 10.1016/j.cell.2013.11.028

Pettersen, E. F., Goddard, T. D., Huang, C. C., Meng, E. C., Couch, G. S., Croll, T. I., Morris, J. H., & Ferrin, T. E. (2021). UCSF ChimeraX: Structure visualization for researchers, educators, and developers. Protein Science : A Publication of the Protein Society, 30(1), 70–82. 10.1002/PRO.3943

Pinto, D., São-José, C., Santos, M. A., & Chambel, L. (2013). Characterization of two resuscitation promoting factors of Listeria monocytogenes. Microbiology (United Kingdom), 159(PART7), 1390–1401. 10.1099/mic.0.067850-0

Popowska, M., & Markiewicz, Z. (2006). Characterization of Listeria monocytogenes protein Lmo0327 with murein hydrolase activity. Archives of Microbiology, 186(1), 69–86. 10.1007/s00203-006-0122-8

Ramijan, K., Ultee, E., Willemse, J., Zhang, Z., Wondergem, J. A. J., van der Meij, A., Heinrich, D., Briegel, A., van Wezel, G. P., & Claessen, D. (2018). Stress-induced formation of cell wall-deficient cells in filamentous actinomycetes. Nature Communications 2018 9:1, 9(1), 1–13. 10.1038/s41467-018-07560-9

Raschle, S., Stephan, R., Stevens, M. J. A., Cernela, N., Zurfluh, K., Muchaamba, F., & Nüesch-Inderbinen, M. (2021). Environmental dissemination of pathogenic Listeria monocytogenes in flowing surface waters in Switzerland. Scientific Reports, 11(1), 9066. 10.1038/S41598-021-88514-Y

Rismondo, J., Haddad, T. F. M., Shen, Y., Loessner, M. J., & Gründling, A. (2020). GtcA is required for LTA glycosylation in Listeria monocytogenes serovar 1/2a and Bacillus subtilis. The Cell Surface, 6, 100038. 10.1016/j.tcsw.2020.100038

Robben, C., Fister, S., Witte, A. K., Schoder, D., Rossmanith, P., & Mester, P. (2018). Induction of the viable but non-culturable state in bacterial pathogens by household cleaners and inorganic salts. Scientific Reports, 8(1), 15132. 10.1038/s41598-018-33595-5

Schardt, J., Jones, G., Müller-Herbst, S., Schauer, K., D’Orazio, S. E. F., & Fuchs, T. M. (2017). Comparison between Listeria sensu stricto and Listeria sensu lato strains identifies novel determinants involved in infection. Scientific Reports, 7(1), 17821. 10.1038/s41598-017-17570-0

Scheinpflug, K., Krylova, O., & Strahl, H. (2017). Measurement of Cell Membrane Fluidity by Laurdan GP: Fluorescence Spectroscopy and Microscopy. Methods in Molecular Biology (Clifton, N.J.), 1520, 159–174. 10.1007/978-1-4939-6634-9_10

Sharma, M., Handy, E. T., East, C. L., Kim, S., Jiang, C., Callahan, M. T., Allard, S. M., Micallef, S., Craighead, S., Anderson-Coughlin, B., Gartley, S., Vanore, A., Kniel, K. E., Haymaker, J., Duncan, R., Foust, D., White, C., Taabodi, M., Hashem, F., … Sapkota, A. R. (2020). Prevalence of Salmonella and Listeria monocytogenes in non-traditional irrigation waters in the Mid-Atlantic United States is affected by water type, season, and recovery method. PloS One, 15(3). 10.1371/JOURNAL.PONE.0229365

Shen, Y., Boulos, S., Sumrall, E., Gerber, B., Julian-Rodero, A., Eugster, M. R., Fieseler, L., Nyström, L., Ebert, M.-O., & Loessner, M. J. (2017). Structural and functional diversity in Listeria cell wall teichoic acids. Journal of Biological Chemistry, 292(43), 17832– 17844. 10.1074/jbc.M117.813964

Signoretto, C., Lleò, M. D. M., & Canepari, P. (2002). Modification of the peptidoglycan of Escherichia coli in the viable but nonculturable state. Current Microbiology, 44(2), 125–131. 10.1007/S00284-001-0062-0/METRICS

Signoretto, C., Lleò, M. D. M., Tafi, M. C., & Canepari, P. (2000). Cell wall chemical composition of Enterococcus faecalis in the viable but nonculturable state. Applied and Environmental Microbiology, 66(5), 1953–1959. 10.1128/AEM.66.5.1953-1959.2000/ASSET/F7488EDF-CD74-4426-A492-E3DC609E8EFE/ASSETS/GRAPHIC/AM0501855001.JPEG

Slavchev, G., Michailova, L., & Markova, N. (2013). Stress-induced L-forms of Mycobacterium bovis: a challenge to survivability. The New Microbiologica, 36(2), 157–166.

Stiefel, P., Schmidt-Emrich, S., Maniura-Weber, K., & Ren, Q. (2015). Critical aspects of using bacterial cell viability assays with the fluorophores SYTO9 and propidium iodide. BMC Microbiology, 15(1), 36. 10.1186/s12866-015-0376-x

Strimmer, K. (2008). fdrtool: A versatile R package for estimating local and tail area-based false discovery rates. Bioinformatics, 24(12), 1461–1462. 10.1093/bioinformatics/btn209

Sun, L., Rogiers, G., Courtin, P., Chapot-Chartier, M.-P., Bierne, H., & Michiels, C. W. (2021). AsnB Mediates Amidation of Meso-Diaminopimelic Acid Residues in the Peptidoglycan of Listeria monocytogenes and Affects Bacterial Surface Properties and Host Cell Invasion. Frontiers in Microbiology, 12(October). 10.3389/fmicb.2021.760253

Talibart, R., Denis, M., Castillo, A., Cappelier, J. M., & Ermel, G. (2000). Survival and recovery of viable but noncultivable forms of Campylobacter in aqueous microcosm. International Journal of Food Microbiology, 55(1–3), 263–267. 10.1016/s0168-1605(00)00201-4

Touche, C., Hamchaoui, S., Quilleré, A., Darsonval, M., & Dubois-Brissonnet, F. (2023). Growth of Listeria monocytogenes is promoted at low temperature when exogenous unsaturated fatty acids are incorporated in its membrane. Food Microbiology, 110, 104170. 10.1016/j.fm.2022.104170

Van Langendonck, N., Bottreau, E., Bailly, S., Tabouret, M., Marly, J., Pardon, P., & Velge, P. (1998). Tissue culture assays using Caco-2 cell line differentiate virulent from non-virulent Listeria monocytogenes strains. Journal of Applied Microbiology, 85(2), 337–346. 10.1046/j.1365-2672.1998.00515.x

Wang, X., Kim, Y., Ma, Q., Hong, S. H., Pokusaeva, K., Sturino, J. M., & Wood, T. K. (2010). Cryptic prophages help bacteria cope with adverse environments. Nature Communications, 1(1), 147. 10.1038/ncomms1146

Weller, D., Wiedmann, M., & Strawn, L. K. (2015). Irrigation Is Significantly Associated with an Increased Prevalence of Listeria monocytogenes in Produce Production Environments in New York State. Journal of Food Protection, 78(6), 1132–1141. 10.4315/0362-028X.JFP-14-584

Whiteley, A. T., Pollock, A. J., & Portnoy, D. A. (2015). The PAMP c-di-AMP Is Essential for Listeria monocytogenes Growth in Rich but Not Minimal Media due to a Toxic Increase in (p)ppGpp. [corrected]. Cell Host & Microbe, 17(6), 788–798. 10.1016/j.chom.2015.05.006

Wideman, N. E., Oliver, J. D., Crandall, P. G., & Jarvis, N. A. (2021). Detection and Potential Virulence of Viable but Non-Culturable (VBNC) Listeria monocytogenes: A Review. Microorganisms, 9(1), 1–11. 10.3390/microorganisms9010194

Witte, C. E., Whiteley, A. T., Burke, T. P., Sauer, J.-D., Portnoy, D. A., & Woodward, J. J. (2013). Cyclic di-AMP is critical for Listeria monocytogenes growth, cell wall homeostasis, and establishment of infection. MBio, 4(3), e00282–13. 10.1128/mBio.00282-13

Wohlfarth, J. C., Feldmüller, M., Schneller, A., Kilcher, S., Burkolter, M., Meile, S., Pilhofer, M., Schuppler, M., & Loessner, M. J. (2023). L-form conversion in Gram-positive bacteria enables escape from phage infection. Nature Microbiology, 8(3), 387–399. 10.1038/s41564-022-01317-3

Yoon, Y., Lee, H., Lee, S., Kim, S., & Choi, K.-H. (2015). Membrane fluidity-related adaptive response mechanisms of foodborne bacterial pathogens under environmental stresses. Food Research International, 72, 25–36. 10.1016/j.foodres.2015.03.016

Zhao, X., Zhong, J., Wei, C., Lin, C.-W., & Ding, T. (2017). Current Perspectives on Viable but Non-culturable State in Foodborne Pathogens. Frontiers in Microbiology, 8(APR), 580. 10.3389/fmicb.2017.00580

